# A Wireless Network of Implantable Microstimulators for Addressable and Patterned Electrical Brain Stimulation

**DOI:** 10.1101/2024.05.25.595909

**Authors:** J. Lee, A. H. Lee, V. Leung, L. Larson, A. Nurmikko

**Author notes:** These authors contributed equally (J. Lee, A. H. Lee).

## Abstract

Transmitting meaningful information into brain circuits by electronic means is a challenge facing brain-computer interfaces. A key goal is to find an approach to inject spatially structured local current stimuli across swaths of sensory areas of the cortex. Here, we introduce a fully wireless approach to multipoint patterned electrical microstimulation by a spatially distributed epicortically implanted network of silicon microchips to target specific areas of the cortex. Each sub-millimeter-sized microchip harvests energy from an external radio-frequency source and converts this into biphasic current injected focally into tissue by a pair of integrated microwires. The amplitude, period, and repetition rate of injected current from each chip are controlled across the implant network by implementing a pre-scheduled, collision-free bitmap wireless communication protocol featuring sub-millisecond latency. As an in vivo demonstration, a network of 30 wireless stimulators was chronically implanted into motor and sensory areas of the cortex in a freely moving rat for three months. We explored the effects of patterned intracortical electrical stimulation on trained animal behavior at average RF powers well below safety limits.

## Introduction

Electrical stimulation of the nervous system has a venerable history. Many important discoveries have led to valuable clinical therapies such as deep brain stimulation to manage Parkinsonian conditions [1, 2, 3], or the use of surface stimulation to identify eloquent areas prior to neurosurgery [4, 5, 6]. In the contemporary field of brain-computer interfaces (BCI), a key aspiration is to advance electronic means to write meaningful information into the cortex. Major advances have been made in *recording* cortical signals which, once successfully decoded, have enabled recent demonstrations of paralyzed people communicating via keyboard commands [7, 8, 9] and translating verbal motor intentions to machine-generated speech [10, 11, 12]. For a future closed-loop BCI system, however, the ability to provide meaningful and naturalistic electrophysiological feedback directly “*into*” the brain in real time remains an elusive goal.

In the case of focused electrical stimulation, there is considerable evidence that current pulses injected locally, e.g., into the hand area of the somatosensory cortex, can elicit tactile sensation in non-human [13, 14, 15, 16] and human primates [17, 18, 19]. Injecting retinotopically matched current stimulus into the primary visual cortex of a non-human primate has also been reported to elicit distinct visual corollaries [20, 21, 22]. In these works, silicon microelectrode arrays (MEAs), such as the “Utah array” [23, 24, 25], have been adopted to deliver simple patterns of current stimulation in assessing the efficacy of microstimulation. However, such monolithic tethered implants have drawbacks in fixed electrode configuration, a finite number of channels, and hence limited cortical coverage. For example, in early clinical trials [26, 27, 28], an MEA evoked sensation only in part of one or two fingers [26, 28]. Scaling up to cover additional cortical areas and implanting multiple MEAs may be impractical for long-term chronic use. Likewise, percutaneous connections are undesirable, highlighting the importance of a wireless implant. We note that delivering focal currents into the cortex has recently been achieved in microprobe designs using magnetic induction as the mechanism [29, 30].

Recently, a wireless multichannel cortical stimulator motivated by prospects for visual prostheses was introduced, which enables the delivery of patterns of current through a monolithic microelectrode array [31, 32]. In this particular embodiment, a coil, an application-specific integrated circuit (ASIC), and a microelectrode array were heterogeneously assembled on a ceramic substrate, resulting in a relatively large form factor for a chronic implant. Thus, there is impetus to engineer a scalable, versatile, and fully wireless miniaturized neurostimulation system which is capable of delivering possibly complex spatiotemporal patterns at a large number of cortical points and across multiple cortical areas.

As an alternative to monolithic multielectrode arrays, autonomous free-standing wireless microchips have been suggested as an avenue toward BCI-related stimulatory interventions [33, 34, 35, 36]. Khalifa et al. reported a miniaturized version of a wireless neural implant based on monolithic silicon integrated circuits aimed at stimulating the cortex or peripheral nerves [34]. However, the design was limited to a single device/channel due to its radiofrequency (RF) communication configuration. In a related scheme, an ultrasound-powered neural microstimulator was introduced for peripheral nerve stimulation [35]. Ultrasound powering allows for deeper penetration into soft tissue than RF; however, the size of the stimulator device remained relatively large because of constraints imposed by the heterogeneous material assembly. In this case, too, only a single device was demonstrated in an anesthetized rodent model. Elsewhere, an endovascular neural stimulator housing a magnetoelectric film has been reported, constrained to a single device so far [36]. A representative summary of contemporary efforts to develop wireless neural microstimulators is given in Supplementary Table 1.

In the context of neural recording, our group previously introduced the idea of spatially distributed autonomous silicon microchips — “neurograins” — as a readily scalable wireless neural *recording* technology, which could be complemented with the capability for intracortical *stimulation* [37, 38, 39, 40]. In this paper, we demonstrate the concept of a scalable multipoint wireless electrical stimulation platform by introducing a novel class of implantable system-on-chip (SoC) silicon microchips. Power to sub-millimeter-sized chips (300 μm to 500 μm in linear dimension) is harvested from an external RF source near 1 GHz and converted into pulsed biphasic currents of up to 120 μA peak-to-peak, deliverable to a specific target cortical area in which each particular chip is implanted.

A key innovation in the paper is the development of a method to program the stimulation parameters (current amplitude, period per phase, and repetition rate) across a large network of chips in real time. We conceived a custom collision-free low duty-cycle communication protocol, a wireless register-mapping method denoted as a ‘daisy-chain’ protocol. Below, we demonstrate the system implemented as a chronic implant in a freely moving rodent. In the communication scheme, the external RF transmitter issues pre-scheduled commands across the chip population via a high-speed downlink at 1 Mbps, addressing and acknowledging the current release events from each device at rates which are instantaneous on a neurophysiological timescale. The approach allows us to control a network of a large number of microdevices and transmit wireless power to generate space-time patterned stimulation only during the delivery of an actual pulsed current stimulus. This significantly reduces the total average RF power exposure, ensuring it remains well below regulatory safety limits, a major advantage of the proposed system.

We assessed the performance of the full wireless system in two *in vivo* phases. Following benchtop validation by injecting currents from a population of microchips into saline, we applied patterns of intracortical microstimulation in anesthetized rats and recorded the evoked neural responses over a range of stimulus conditions. We then implanted 30 microchips across the cortex of a rat for chronic experiments and applied spatially targeted stimulation while selectively sampling the vast stimulus parameter space in a freely moving animal over three months. Distinct neuromodulation effects were measured as the animal engaged in a trained lever pressing task. Lastly, although the size of the rat brain limits the number of implants, extrapolation of the experimental data suggests that a wireless system composed of up to 1000 microstimulators can be accessed in less than 3 milliseconds to deliver complex dynamical patterns of cortical excitation.

### Overview of Implantable Wireless Microstimulators – Circuits and Communication

Fig. 1a illustrates the principle of the multipoint wireless microstimulator implant for chronic experiments in a rat model. A population of spatially distributed autonomous microchips is deposited on the cortex of the animal and energized from epidermal and subdermal RF coils by near-field inductive coupling. Two coils are specifically designed to match the rat’s cortical geometry, covering an area of 8 mm × 13 mm on the cortex. Simultaneously, the downlink also transmits stimulus-specific instructions to the network. For delivery of intracortical currents, pairs of tungsten microwires separated by 100 μm were affixed to the epicortical chips in a post-process step (Fig. 1b, Supplementary Figure 1). An example of an ensemble of spatially distributed functional stimulator chips embedded in an agarose brain phantom is shown in the photo of Fig. 1c to illustrate the autonomy of their individual placement.

**Figure 1.**
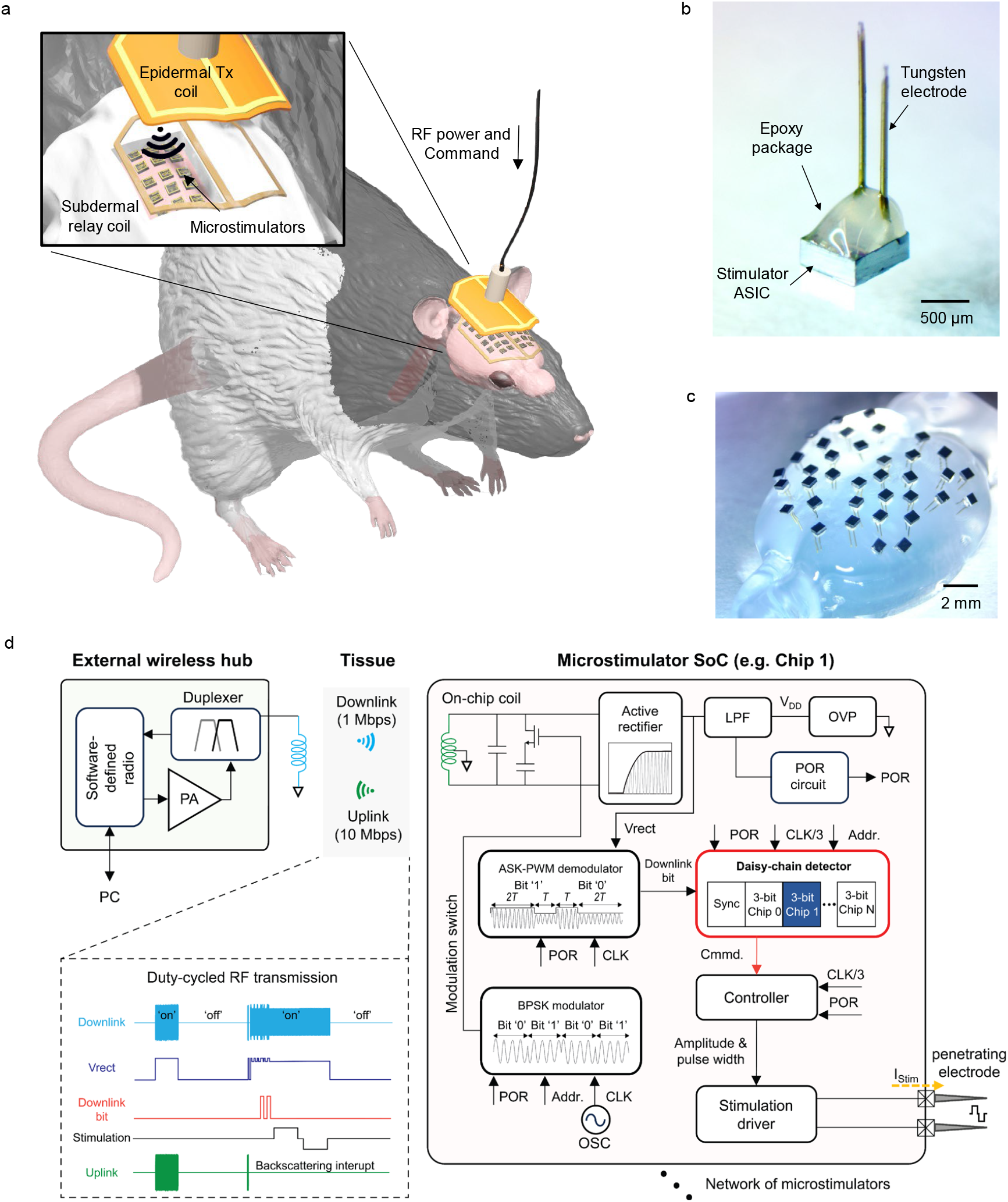
Distributed microstimulator system. **a**. Schematic showing a population of microchips implanted on rat cortex together with epi- and subdermal coils for resonant near-field RF energy transfer and delivery of stimulation commands. **b**. Microphotograph showing a stimulator ASIC with integrated tungsten electrodes to enable intracortical microstimulation. **c**. Photo of microchips inserted into a brain phantom made of agar to test the distributed wireless interface on benchtop. **d**. Design of microchip circuitry and RF daisy-chain communication scheme for low-duty cycle operation. The downlink delivers RF power and commands at 1 Mbps to specific microchips to select one of preprogrammed current waveforms. The ASIC consists of power harvesting, data recovery, daisy-chain detector, controller, and stimulation circuit blocks. Power-on-reset triggers a BPSK modulation circuit to report its unique address, driven by a free-running oscillator. The inset shows the waveform of the downlink and uplink, rectifier output voltage (Vrect), demodulated downlink bit, and stimulation. The plot illustrates how the downlink is ‘on’ for a short period of time to deliver RF energy to the chip – only when needed resulting in duty-cycled RF transmission hence lowering average RF exposure. Abbreviations: V_DD_: voltage supply, OVP: over voltage protection, POR: power-on reset, LPF: low pass filter, OSC: oscillator, CLK: clock, Addr.: address, Cmmd.: command, PA: power amplifier, PC: personal computer, BPSK: binary phase shift keying, Vrect: rectified downlink signal.

Fig. 1d shows the transcutaneous RF link (downlink and uplink), a schematic of the timing diagram, and a block circuit description of an individual microchip. The circuits employ the daisy-chain, register-mapping communication scheme designed for speed and scalability to accommodate a large number of chips with minimum aggregate latency (Supplementary Note 1). A three-bit downlink instruction delivered in mere 3 μs programs a specific stimulation waveform for an individual chip; hence, in principle, arbitrary patterns of microstimulation can be generated by the daisy-chain scheme for up to a thousand chips within 3 ms. External supporting RF transmitter electronics in the experiments included a software-defined radio (SDR) to generate the downlink commands and collect the uplink signals, an RF power amplifier, and miscellaneous microwave components.

As described next, the integrated circuit of a current injecting microchip contains an amplitude shift keying pulse width modulation (ASK-PWM) demodulator for decoding downlink signals, a daisy-chain detector, a digital controller, a current source driver (see details in Supplementary Figure 2), a free-running oscillator, and a binary phase shift keying (BPSK) modulator for backscattering (uplink) whose purpose is to report the status of each chip to the transmitter hub. The envelope of the rectified incoming RF signal is interpreted by the custom ASK-PWM demodulator, which is capable of downlink bit recovery tolerant to a variance of the free-running oscillator frequency (Supplementary Note 2) [37, 40].

The ‘daisy-chain detector’ (in the red rectangle in the ‘Microstimulator box’ of Fig. 1d) monitors downlink bits as generated by the demodulator. Upon detecting the proper ‘sync’ sequence, the detector begins counting bits until finding the three bits assigned to the specific chip as identified by its unique address (Supplementary Note 3). Then, the digital controller uses these three bits to program the specific stimulation waveform for the current source driver. Upon chip activation (turn-on), each microchip generates an uplink signal through RF backscattering for 96 μs, triggered by a power-on reset (POR), which reports its unique on-chip address (see Supplementary Note 4). Note that, as the clock frequency of a chip can be extracted from the received binary phase-shift keying (BPSK) uplink data, these data can be used as an indicator for the power status onboard a chip.

### Programming and Near-field RF Powering Microstimulator Ensembles

To ensure safe RF power delivery across ensembles of spatially distributed microchips, we considered the likely variations in the magnetic field across the total implant area and the constraints on energy harvesting. We implemented the stimulator circuit of Fig. 1d in silicon dies with sizes of 300, 400, and 500 μm, fabricated in the 65 nm low-power RF CMOS process. We measured how the size of the microcoil on the die affects the incident transmitting (Tx) power requirements (Fig. 2a). To maximize the available on-chip charge, we used an unregulated voltage supply for the overall circuit where an overvoltage protection (OVP) diode sets the upper limit of V_DD_. Thus, the RF power affects the harvested power level as well as the voltage supply so that an increase in RF power causes an increase in the clock frequency of the free-running on-chip oscillator. This test also suggests that even smaller microstimulators can operate in a similar fashion albeit with increased RF power. In the animal experiments, we primarily used the slightly larger 500 μm size chips due to the availability of larger numbers of chips and higher fabrication yield in microwire integration.

**Figure 2.**
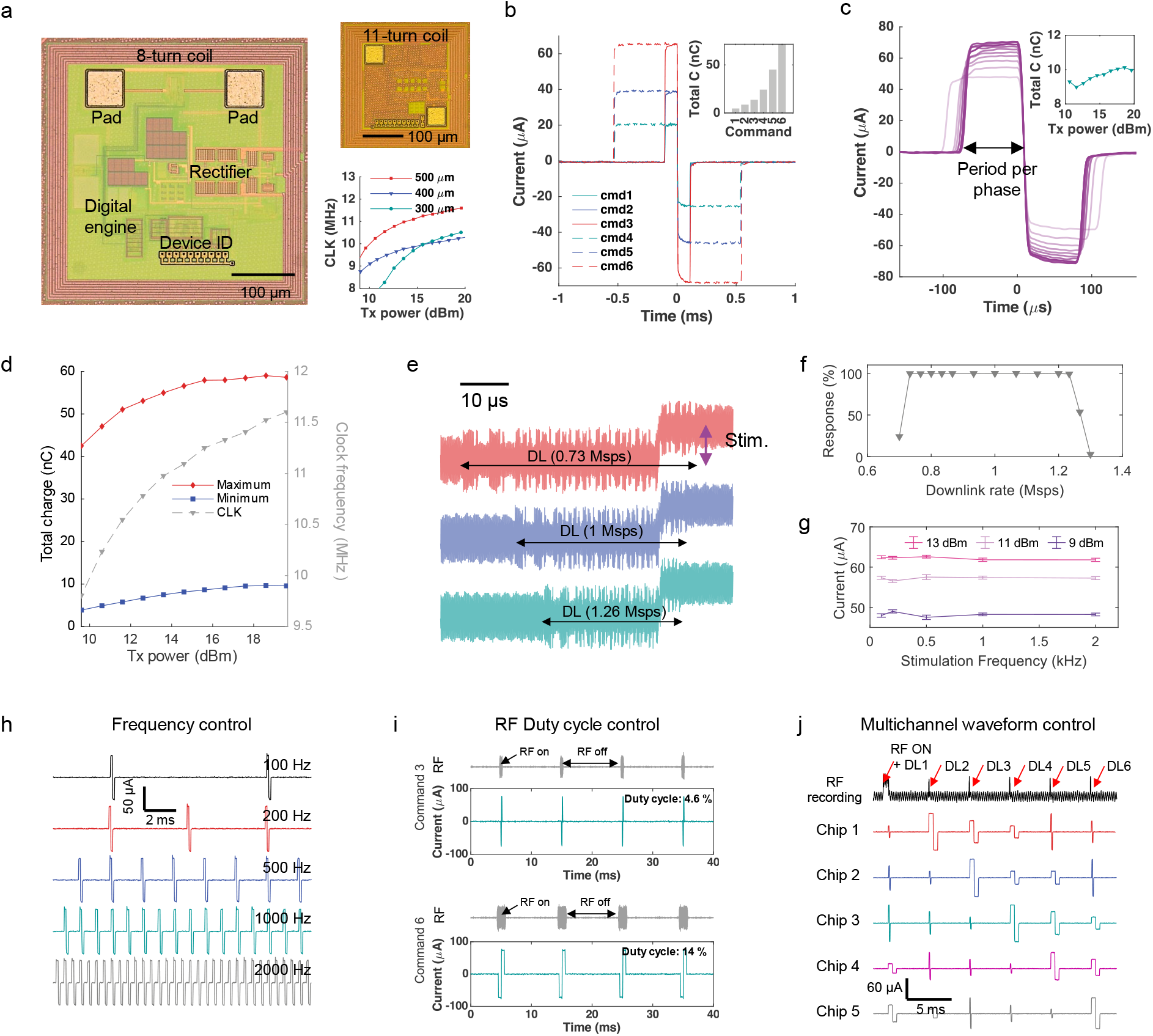
Properties of microstimulator ASICs. **a**. Microphotograph of two stimulator chips of 400 μm × 400 μm and 300 μm × 300 μm size, respectively. The inset shows how on-chip clock frequency depends on the incident Tx power (benchtop test). **b**. Current waveforms generated by six distinct downlink commands to vary both the amplitude and pulse width. The inset shows the total injected charge per specific command. **c**. Expanded time axis of the stimulation waveform for a 200 μs pulse width (100 μs per phase) at varying Tx power levels. The inset shows the total injected charge as a function of the Tx power. **d**. Dependence on Tx power level for a maximum (Command 6) and a minimum (Command 1) of charge injection and on-chip clock frequency, respectively. **e**. Downlink (DL) waveforms transmitted at different data rates (0.73-1.26 Msps) and the stimulation event generated as a response to the commands for chip address 6. **f**. Tolerance to a wide range of downlink rates when measured as a response (stimulation) to downlink commands across 1000 trials. **g**. The amplitude of current injection as a function of Tx RF power and a wide range of stimulation frequencies, demonstrating stable current injection up to 2 kHz (100 μs per phase). Data points present average values, and error whiskers show average ± standard deviation (n=100). **h**. Control of the stimulation frequency directly by downlink transmission frequency. **i**. Waveforms of the RF downlink and injected stimulation, respectively, demonstrating low-duty cycle RF transmission at 100 Hz frequency. The duty cycle is lower for stimulation of 100 μs per phase (top, Command 3) case, compared to 500 μs per phase (bottom, Command 6). **j**. Programmability of temporal stimulation across a five-chip population; a single downlink sequence determines the stimulation amplitude and period per phase simultaneously for all chips. In these experiments, the stimulation was delivered into 14 Kohm load resistance. Abbreviations: ID: identification, CLK: clock, cmd: command, C: charge, stim.: stimulation.

Three-bit commands from the downlink (seven types, 6 types for single shot stimulation, 1 type for continuous 100 Hz stimulation, Supplementary Table 2) determine the current amplitude and pulse period of injected current as shown in Fig. 2b, where each command corresponds to the delivery of a specified amount of total charge (Fig. 2b inset). Due to the unregulated voltage supply, the current amplitude increases with respect to the incoming RF power as shown in Fig. 2c. However, since the clock frequency also increases with RF power, the stimulation period decreases. As a result, the two effects nearly cancel out so that the increase in injected charge (e.g. for Command 3) is up by only 13.7% despite a tenfold increase in incident Tx power (the inset of Fig. 2c). As an example, Fig. 2d illustrates the total injected charge generated by Commands 6 (maximum) and 1 (minimum), respectively. Note that, in this case, if the Tx power were to increase tenfold, the increase in delivered maximum charge was measured as only 38.8%, demonstrating the moderation of RF power level variations in the circuit design notwithstanding the choice of the simple unregulated voltage source.

The free-running oscillator in the microstimulator offers the benefit of low power consumption and a small footprint. Still, wafer-level variations in semiconductor fabrication can lead to clock frequency variance across an ensemble of RF-powered devices [37, 40]. Our ASK-PWM downlink protocol (Supplementary Note 2) is tolerant to anticipated clock variance; a feature validated by varying the downlink data rate as shown in Figs. 2e and 2f. The results demonstrate that the ASK-PWM demodulator can achieve a 100% success rate provided that the downlink clock variance is less than 23%, a value well within the range of observed clock frequency variance in our chips. The downlink protocol also allows the generation of a wide range of stimulation frequencies, here up to 2 kHz even if such frequencies might be well in excess of typical pulse repetition rates for neuromodulation. Even for such high-frequency stimulation, however, the amplitude of the injected pulsed current remains rather constant as shown in Figs. 2g and 2h where only the incident Tx power determines the amplitude of the current.

Since the period of active current injection in most neuromodulation schemes is relatively short, on the order of 100 μs [26, 27], the wireless RF energy transfer can likewise be kept short to reduce the average RF power low. Fig. 2i demonstrates such low-duty cycle RF transmission by displaying the 100 Hz biphasic stimulation waveform with 100 μs and 500 μs per phase generated by a downlink at 4.6% and 14% duty cycle, respectively. Crucially, such a short period downlink command can control multiple devices simultaneously and reliably, as demonstrated by benchtop measurements in Fig. 2j with five chips. In this example, six different downlink commands are transmitted sequentially, each command generating preprogrammed values of current amplitudes and periods per phase across the set.

### Intracortical Microstimulation on Benchtop and in Acute In Vivo Rat Model

After attaching a pair of tungsten microwires (Fig. 1b) onto each ASIC die for focal intracortical current injection (see Supplementary Figure 1 for microwire post-process integration procedure), we first measured the pulsed current injection on the benchtop both by individual as well as a population of 24 microstimulators. Using phosphate-buffered saline (PBS) as a substitute for conducting tissue as illustrated in Fig. 3a, we first measured the RF-powered injected current using a sensing resistor in series as shown in Supplementary Figure 3. An example of both the biphasic currents and applied voltages generated by the ASIC is shown in the top row traces of Fig. 3b where the pulse shapes include finite effects from the capacitive interface between the tungsten electrodes and the saline electrolyte. According to each specific wireless command, the chip delivers varying amounts of charge as seen in the bottom row data of Fig. 3b. While a finite charge imbalance remains, this amount is small relative to the total injected charge (generally less than 2%) because of active charge balancing. We then used a custom (wired) 24-channel planar recording electrode (inset of Fig. 3a), to illustrate the generation of arbitrary spatiotemporal patterns by a 24-element wireless network of microstimulators, here dynamically writing out the sequence of a series of letters spelling ‘BROWN’ in Fig. 3c.

**Figure 3.**
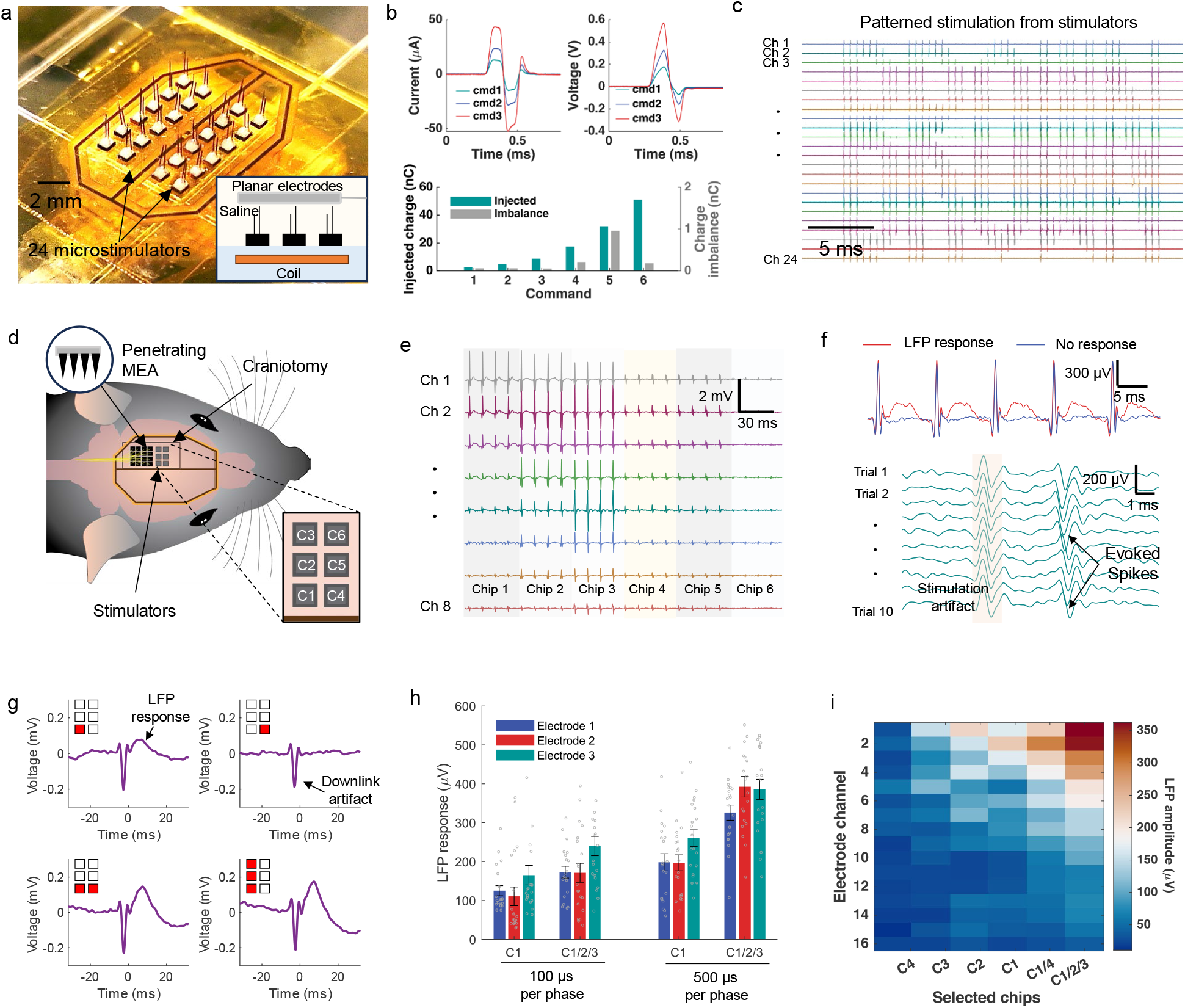
Demonstration of pattern stimulation by the wireless stimulator network on benchtop and in-vivo acute experiments. **a**. Immersion of microchips in transparent saline. The inset shows the location of the relay coil (bottom of the dish) and a 24-element planar recording array with its size matching the coil underneath. **b**. Voltage and current waveforms of stimulation injected by the microstimulator into the saline through a pair of tungsten electrodes (top row). The bottom row shows injected total charge and charge imbalance depending on the stimulation waveforms (all plots averaged over 60 trials). **c**. Spatiotemporal stimulation waveforms generated by 24 microchips to inject a current pattern spelling “BROWN”. **d**. Microchips implanted into a rat cortex in acute experiments (six chips C1 to C6, and the 16 channels penetrating MEA, separation of adjacent silicon needles is 400 μm). **e**. A plot generated by selecting each of the 6 stimulators and injecting current, captured by 8 recording channels in MEA as the ‘stimulation artifact’ (Command 7). **f**. LFP responses evoked by microstimulation (red, top) compared to the absence of response observed through the same electrode when repeated stimulation led to apparent neuronal fatigue (blue, top). The plot at the bottom displays the evoked neuronal spike responses following each stimulation event. **g**. Averaged LFP responses and downlink artifacts caused by rapid RF power switching during duty-cycled RF transmission, here for chips delivering currents with 100 μs period per phase (Command 3, 10 trials, bandpass-filtered from 60 to 2000 Hz). **h**. Example showing the dependence of LFP response amplitudes on the stimulation period and the specific set of chips selected for stimulation. Bars present average values and error whiskers show average ± standard error (n=20, dots for individual data points). **i**. Composite heat map of LFP response amplitudes induced by selected chips (500 μs period per phase, Command 6) and measured across 16 recording electrodes to demonstrate both local and broad cortical modulation (averaged over 20 trials). Maximum Tx power of 20 dBm (100 mW) was applied in the acute animal experiments.

We next implanted six chips into the cortex of a rat to assess the in vivo wireless stimulation capability while the animal was under anesthesia. Given the large parameter space in terms of all possible combinations of stimulation patterns of variable current amplitude, pulse width, and frequency, we did not aim in this paper at a detailed neuroscience study. Rather, we aimed to test the functionality of the multipoint wireless microchip system by selective sampling of the multidimensional stimulus parameter space. A wired ‘Utah’-type multielectrode array (penetrating MEA, channels 1-16) was implanted adjacent to the wireless stimulating microchips (Fig. 3d) to monitor both the current waveforms in cortical tissue and to record the waveforms of evoked neural responses. Given that the distance from each current injecting microchip relative to a given MEA recording electrode varied, we could collect the matrix of data in Fig. 3e to verify that the wirelessly controlled address selection works reliably for an in vivo implant.

We probed the effect of focal microstimulation in selected locations of the cortex while actuating single or multiple microchips, achieving both local and broad cortical modulation. For example, when stimulating the cortex at 100 Hz using a given single microchip, we recorded both evoked local field potentials (LFPs) and extracellular spike activities (Fig. 3f). As shown in Fig. 3g, current simultaneously injected by chips 1, 2, and 3 evoked higher amplitude LFP responses compared to that by single chip 1 alone. In addition, a longer period stimulation (500 μs current pulse vs. 100 μs pulse) evoked larger LFPs presumably because of the increased total injected charge as shown in Fig. 3h [41]. The results of the paired t-test analysis comparing conditions in Fig. 3h are summarized in Supplementary Table 3, which shows that both the multichip stimulation and the period of stimulation have statistically significant effects. These results also suggest that an equivalent level of cortical neuromodulation can be evoked by short-period stimulation delivered by multiple chips when compared to a longer stimulus by a single chip. Note that the former case corresponds to a lower average applied RF power due to the lower duty cycle. Furthermore, temporally simultaneous stimulation from multiple chips generated robust LFP responses across a wider cortical area as illustrated in Fig. 3i and Supplementary Figure 4. Here, stimulating the cortex by chips 1, 2, and 3, the recordings from the MEA displayed distributed LFP responses larger than 100 μV at up to ten electrode sites whereas only five MEA electrodes could record comparable LFP responses from stimulation by chip number 1 alone.

**Figure 4.**
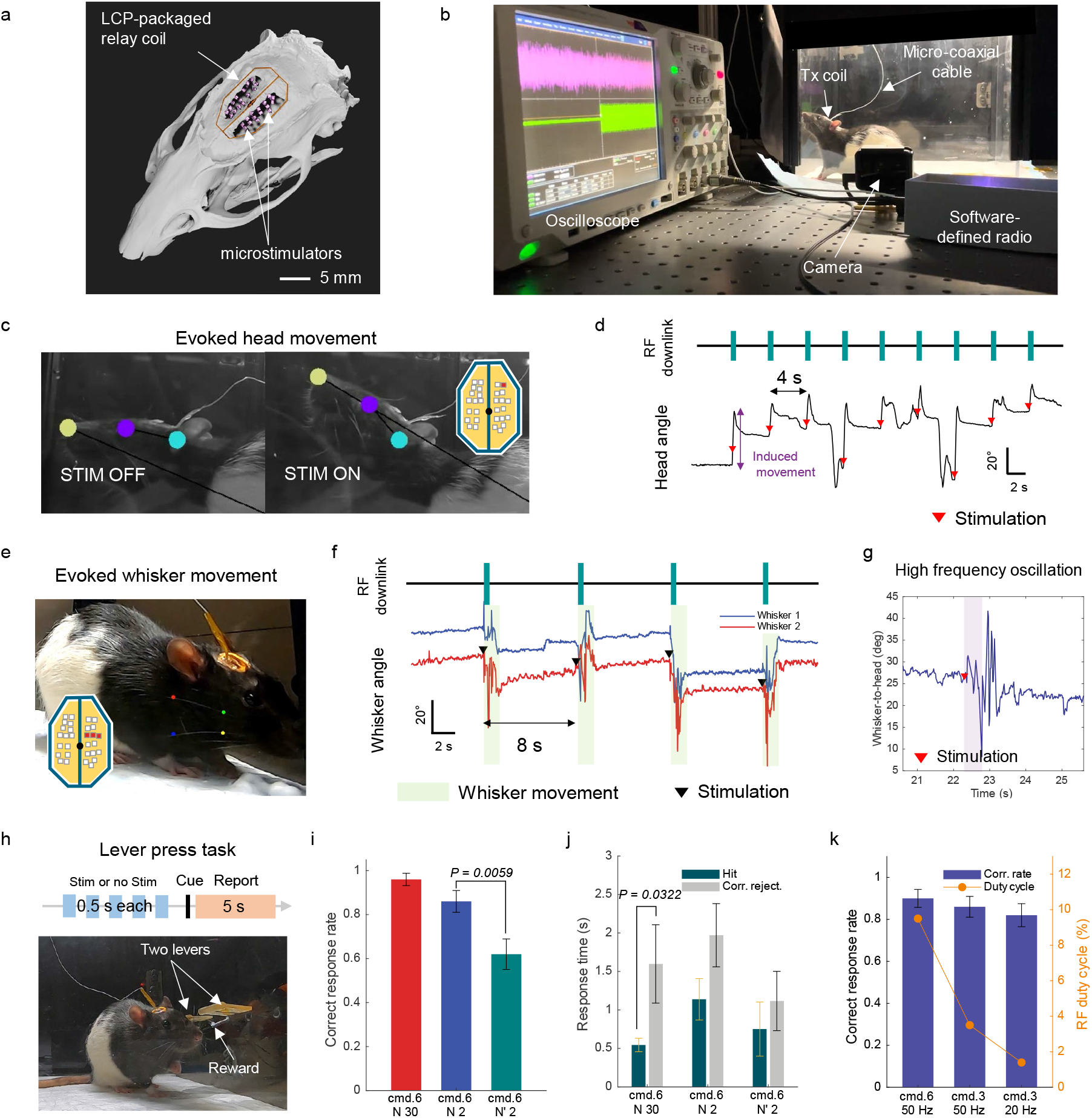
Microchip implants in freely moving chronic rat model. **a**. Postmortem micro-CT scan image of the animal’s head showing the location of the relay coil and microstimulators (additional views in Supplementary Figure 5). **b**. The experimental setup includes a custom enclosure allowing the rat to move freely while the Tx coil atop the head; supporting instrumentation includes a software-defined radio (SDR), an oscilloscope to monitor the downlink, and a camera. **c**. Captured video images showing the head position before and following stimulation. The inset shows the location of the particular microchip (in red) from a population of 30 chips delivering stimulation (400 Hz, 250 ms stimulation pulse train duration, and 500 μs per phase, the black dot in the center is aligned with the bregma). **d**. Timing and duration of the RF downlink to trigger focal microstimulation of the motor cortex and measured head angles relative to the horizon after each stimulation event. **e**. Photo of animal and the two target whiskers: whisker 1 labeled with red and green dots, and whisker 2 labeled with blue and yellow dots. The inset shows three microchips (each injecting pulsed current at 400 Hz, 500 μs per phase, and 500 ms train duration). **f**. Timing of the RF commands and stimulus-evoked whisker movement; whisker angles are measured with respect to the horizon. **g**. Post-stimulus, high-frequency oscillatory motion of evoked whisker response. The purple-shaded area on the graph represents the duration of the pulse train. **h**. The timing diagram for the choice lever-pressing task and the lever setup. If the animal perceives the current stimulus, it should report the percept by pressing the left lever; otherwise, it should press the right lever. An audio cue marked the start of a report session. **i**. Correct response rate based on the number of microstimulators delivering current stimulus (the location of N2, N’2, and N1 is shown in Supplementary Figure 9, the trial number is 50, stimulus parameters: 50 Hz, 500 ms pulse train duration, 5 train repeats). **j**. Response (reaction) time for ‘Hit’ and ‘Correct rejection’ in relation to audio cue, respectively. **k**. Dependence of correct response rate and RF duty cycle on the stimulation frequency and the period of each stimulation phase (single chip stimulation, N1, 500 ms train duration, 5 train repeats). In **i, j** and **k**, Bars present average values and error whiskers show average ± standard error (n=50). The p-values shown in the plots were obtained from paired t-tests indicating significant differences (p < 0.05). The Tx power was 20.15 dBm in all chronic experiments. Abbreviations: LCP: liquid crystal polymer, Corr. reject.: correct rejection, Corr: correct.

### Multipoint Intracortical Microstimulation in Chronic In-vivo Rat Model

For a demonstration of spatially distributed intracortical electrical microstimulation in an awake, freely moving animal, we implanted 30 microchips on the surface of the rat cortex together with a resonant RF energy-transferring subcutaneous relay coil (Fig. 1a). The chips were distributed across the surface of motor and sensory areas, each chip equipped with a cortex-penetrating microwire electrode pair. The location and distribution of the chip population were later imaged by postmortem micro-computed tomography (CT) scans acquired more than three months after surgery (Fig. 4a and Supplementary Figure 5). The external Tx coil was mounted onto the head of the animal enabling the rat to move freely within a 33 cm × 33 cm × 42 cm sized enclosure which was instrumented for a behavioral lever-pressing task (Fig. 4b and Supplementary Figure 6). The lightweight Tx coil (170 mg) was connected via a thin microcoaxial cable to an RF duplexer, an amplifier, and the SDR unit for RF power delivery and command/communication. Details of the hardware and the behavioral test setup can be found in Supplementary Figures 7 and 8 and Methods.

We first targeted the right hemisphere of the motor cortex and delivered current stimulus while recording any overt animal responses with a video camera. The current injection of 120 μA (peak-to-peak) by a single microdevice was sufficient to evoke clear head movement as shown in Fig. 4c and Supplementary Video 1. We used high frequency burst stimulation at 400 Hz (500 μs per phase, 0.25-second pulse train duration), fast on the scale of a neuron’s response time but a rate which has been shown to evoke robust muscle twitch in rodents in other intracortical work [42, 43]. ‘pulse train duration’ refers to the time period during which the 400 Hz stimulation was delivered, resulting in a total of 100 current stimulation events in this case. The inset in Fig. 4c shows the location of the particular chip in the motor cortex and its position relative to the perimeter of the relay coil. Fig. 4d plots the measured head angle changes during stimulation events, showing the stimulation evoked a tilt in the head angle and the subsequent corrective/reactive action by the animal then decreased the head angle over several seconds.

We then selected a set of three chips distributed across both the motor and sensory cortex (Fig. 4e). High frequency stimulation (400 Hz, 500 ms pulse train duration, and 500 μs per phase) evoked readily measurable high-frequency oscillatory motion of whiskers, as seen in Supplementary Video 2; the evoked response was also measured as post-stimulus changes in the whisker angle (Fig. 4f). An expanded view of high-frequency oscillations of whisker movement is shown in Fig. 4g. The result suggests that spatial summation of local currents injected from the three chips stimulated the animal’s sensorimotor whisker circuit and triggered a corresponding motor response. To allow sufficient time for the cortical network to reset between stimulus events and to reduce the average RF power in each trial, we reduced the effective duty cycle to 4.37% for both head and whisker movements by incorporating resting periods of 4 seconds and 8 seconds, respectively.

Further exploring the neuromodulation capabilities of the microchip implant, we designed a behavioral experiment where we trained the animal in a two-lever pressing task. The aim was for the animal to report percepts evoked by current stimuli (Fig. 4h). If the rat detects a stimulus, it is expected to press the left lever; if it does not detect the stimulus, it has been trained to press the right lever in both cases to receive a reward (Supplementary Video 3). A sound cue was played to initiate the report. When stimulating all thirty chips simultaneously at 50 Hz frequency, the animal successfully performed the detection task to achieve a 96% rate for the correct lever pressing response. When only two chips from the set were chosen — stimulating the sensory cortex (N2) or motor cortex (N’2) (see positions in Supplementary Figure 9) — the correct response rates were reduced to 86% and 62%, respectively (Fig. 4i). The difference suggests that the animal’s experiential sensation, that is its perception of the stimulus, depended on the cortical location where the focal current was injected. The kinematic response times, i.e. animal’s reaction times for ‘hits’ and ‘correct rejections’ are shown in Fig. 4j. There was a statistically significant difference in the response times for ‘hits’ and ‘correct rejections’ when the 30 microstimulator was actuated, perhaps indicating that the time between the animal’s perception and subsequent reporting depended on the strength of the sensation induced by the current stimulus.

Finally, we tested whether the animal could perceive stimulation delivered at a low duty cycle RF transmission —a key feature of our system, even when delivered by a single microstimulator within the set. In this experiment, we measured the correct response rate in the detection task when stimuli were delivered at two frequencies, 50 Hz and 20 Hz, and two pulse periods per phase, 100 μs and 500 μs (Commands 3 and 6). As shown in Fig. 4k, with a stimulation frequency of 50 Hz and a phase period of 500 μs, the corresponding RF duty cycle was only 9.5%, yet the animal still achieved a 90% correct response rate. Even when the stimulation frequency was reduced to 20 Hz and the period to 100 μs per phase, resulting in an RF duty cycle of only 1.4%, the animal managed an 82% correct response rate. This suggests that even modest charge injection under low RF duty-cycled transmission can evoke perceptual experiences detectable behaviorally. These in-vivo experimental results demonstrate the versatility of our wireless system, which opens the doors to seeking for optimal stimulation strategy for neuromodulation through parameters such as patterned channel selection, current stimulation period, amplitude, and frequency while minimizing RF energy exposure.

### Stability of the Wireless Link in Chronic Implants in Freely Moving Rat

The RF powering of and backscattered signals from the chronic wireless implant involved inductive coupling via the wireless 3-coil system of Fig. 1a, namely the Tx coil above the animal’s skin, the subcutaneous relay coil above the skull, and the chip microcoils located on the cortical surface. Neither the subcutaneous relay coil nor the chips were visible once the scalp was fully healed after 21 days post-surgery as shown in the photo of Fig. 5a. For hermeticity, the subcutaneous relay coil was encapsulated by a 225 μm thick layer of LCP. Its resonant coupling characteristics could be measured by monitoring the relevant S-parameter of the external Tx coil, reflectance S_11_, and a network analyzer. The value of S_11_ as a function of time is plotted in Fig. 5b to illustrate how the relay coil maintained a sharp resonance peak even 70 days after implantation (measurement setup in Supplementary Figure 10).

**Figure 5.**
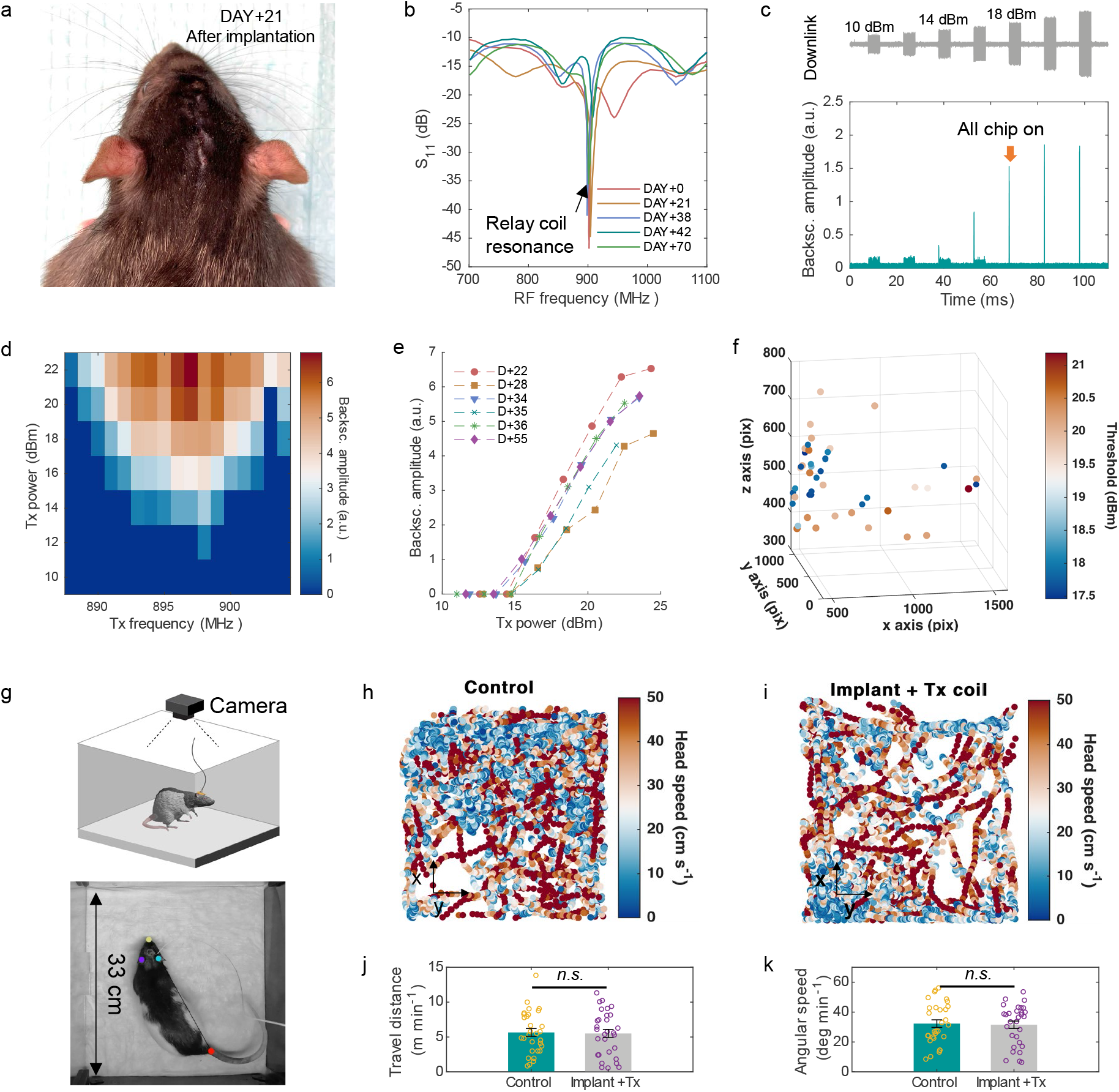
Stability of the 3-coil wireless link in the chronic implant. **a**. A photograph of the animal’s head 21 days after the implantation of the relay coil and 30 microchips. **b**. Measured S-parameter (S_11_) of the Tx coil attached to the head, showing the resonance of the implanted relay coil over two months. **c**. RF downlink signal with varying power amplitudes (top) and amplitudes of backscattered signals from microchip population displaying a threshold Tx power of 18 dBm (bottom). **d**. Backscattered signal amplitudes as a function of Tx power level and Tx frequency. **e**. The amplitude of backscattered signals recorded up to 55 days following implantation and its dependence on Tx power. **f**. Selected x-y locations of the animal freely moving in the enclosure with corresponding threshold Tx power (additional coordinate plane views in Supplementary Figure 11 and video capture examples in Supplementary Figure 12). **g**. Schematic and photo captured by camera used for motion tracking inside the 33 cm × 33 cm × 42 cm sized enclosure. **h** and **i**. The location and speed of the animal’s head tracked for 30 minutes for both a control rat and the rat with implants and the Tx coil attached to its head, respectively. **j** and **k**. Distance traveled per minute and head angular speed per minute, both averaged over 1 minute, showing no statistically significant difference between the control rat and the rat with implants. Bars present average values and error whiskers show average ± standard error (n=30, p-value 0.85 and 0.83 for distance traveled and angular speed, respectively). The p-values were obtained from paired t-tests indicating no significant differences (p > 0.05). Circles represent each data point. Backsc.: backscattering, a.u.: arbitrary unit, n.s.: not significant.

We also examined the strength of the inductive coupling across the microchip population by varying the Tx power and measuring the amplitude of backscattered signals. When all chips were activated, the backscattered signals summed up to a short “reporting” pulse in Fig. 5c. Here, the total amplitude of the backscattered signal is proportional to the number of active devices and the Tx power, but is also affected by variations in the efficiency of the wireless link. We evaluated the effective efficiency of the 3-coil resonant system by varying the Tx frequency and measuring the backscattering amplitude where Fig. 5d shows the resonance at 898 MHz. We monitored the amplitude of the backscattered signal over two months and observed good stability of energy harvesting over time in the freely moving animal, as shown in Fig. 5e. Along with Fig. 5b, these results also indicate that LCP packaging is impermeable to water or electrolyte intrusion at least over two months where such intrusion can lower the quality factor of the coil and/or cause a shift in the resonance frequency.

To test the possible variation in threshold RF power required to turn on all chips while the animal moved inside the 33 cm × 33 cm × 42 cm sized enclosure (Fig 5g), we monitored backscattering signals at multiple x-y coordinate points. The results in Fig. 5f and Supplementary Figure 11 show no measurable dependency of the threshold on the animal’s x-y position. Rather, a more important factor was the variation in its behavior which includes scratching, standing, exploring, and nesting (video capture shots in Supplementary Figure 12). When the animal was still, the average threshold Tx power was 17.78 dBm. Those behaviors that involved direct contact of paws with the head and Tx coil caused the threshold to increase by up to 3.4 dBm. In this worst-case scenario, the misalignment of the coils led to a 2.1-fold increase in the required RF power. Even so, the average power values still remained well below the specific absorption rate (SAR) safety limit.

Video monitoring (Fig. 5g) also allowed us to answer two additional questions: 1) whether the multichip implant and the subdermal relay coil themselves might impair the motor activity of the animal, and 2) whether the microcoaxial cable from the Tx coil to RF electronics hindered its freedom of movement. Figs. 5h and 5i show that the animal was able to move across the enclosure comparably to another animal without the implant (referred to as ‘Control’). Furthermore, the differences in distance traveled per minute or head angular speed per minute between the two sets of data acquired from each animal were not statistically significant, as shown in Figs. 5j and 5k.

## Discussion

In this paper, we have demonstrated the delivery of patterned electrical brain stimulation by a spatially distributed wireless network of implanted silicon microchips. The network can generate arbitrary space-time patterns of stimuli across multiple cortical areas as a chronic rodent implant. One key feature in the design of the wireless system is the RF command-and-control communication protocol which enables virtually instantaneous control of a population of microstimulators to generate complex patterns of current injection. Such spatiotemporal programming is possible under low-duty cycle RF conditions so that the average RF exposure to tissue remains low in a long-term implant. For a proof-of-concept demonstration, we implanted microchips in both anesthetized and freely moving rats to demonstrate evoked neural and behavioral responses to multipoint cortical neuromodulation. The wireless system was able to trigger head and whisker movements by a direct current stimulus into the motor cortex and reliably generate percepts by activating the underlying circuits in the sensory cortex while the animal was engaged in a trained behavioral task.

In the chronic studies, we delivered up to 103.5 mW of peak Tx power directly into the animal’s head. Supplementary Figure 13 shows that the 3-coil system designed here for rodents has a peak spatial SAR value of 1 W/kg (averaged over 10 g of tissue at continuous 100 mW Tx power), an order of magnitude below the tissue safety threshold for human use of 10 W/kg [37, 44, 45]. Yet, by lowering the operational duty cycle to 1.4%, the average RF power required to elicit a behavioral response was a mere 1.45 mW. This excitation was sufficient to achieve an 82% success rate in the correct stimulus response in a lever-pressing task, even by a single microstimulator chip. Therefore, operating in low-duty cycle transmission mode yet providing effective neuromodulation, the corresponding SAR value could be reduced by another order of magnitude down to 0.0145 W/kg for a stimulation period per phase of 100 μs and stimulation frequency 20 Hz (again 1.4% duty cycle). Such stimulus parameters are well within the range found in many electrical neuromodulation experiments in primates [46, 47]. Therefore, even allowing for a tenfold decrease in the efficiency of near-field energy transfer for thicker tissue in a primate (scalp, skull, dura) [37], a SAR value well below even the ‘low-tier’ threshold of 2 W/kg appears to be possible [45].

The wireless technology introduced in this paper offers multiple opportunities for basic neuroscience studies and possibilities for clinical neurotechnologies. The ability to deliver arbitrary spatiotemporal patterns of current stimulus over a wide parameter range with minimal latency can be particularly valuable in a closed-loop BCI system designed for the restoration of sensations [48, 49]. Likewise, the system could offer an alternative approach to the development of visual cortical prostheses with implanted microchips across the primary visual cortex generating phosphene patterns with input from external cameras [50]. As such, our wireless microchip approach is quite general and potentially broadly applicable to biomedical engineering applications which can benefit from multipoint, patterned electrical stimulation of specific physiological targets in the body. These might include cardiac stimulators as multipoint pacemakers designed to correct specific circuit disorders and subdermal microchips to convey tactile feedback to amputees wearing prostheses, to name but two examples.

## Methods

### Microstimulator design and fabrication

We designed the microstimulator ASIC in TSMC’s 65 nm mixed-signal/RF low-power CMOS process. The daisy-chain detector designed for low duty-cycled RF transmission provides a core distinction from the previous design using on-chip address-based call-and-response communication presented in the previous work [37]. After the chip had been fabricated, we integrated a pair of 50 μm diameter tungsten microelectrodes for intracortical access as described in Supplementary Figure 1. We used silver epoxy (H20E, Epo-Tek) to first fabricate ball supports at the edge of electrodes with the curing temperature of 200 °C for 15 min. Subsequently, we placed the electrodes on the chip after applying additional silver epoxy to the 60 μm chip pads. Ensuring perpendicular alignment of the electrodes with the chip surface, we cured the epoxy at a low temperature of 80 °C for 8 hours to prevent chip damage. For mechanical reinforcement and encapsulation, we applied biomedical-grade epoxy (M-31CL, Loctite) to the base of the electrode and the chip. Lastly, the tips of stimulator electrodes were inserted into the polydimethylsiloxane (PDMS) polymer and then the other parts were coated with parylene-C to achieve better biocompatibility and further encapsulation. Chemical vapor deposition formed a 3 μm conformal coating of parylene-C on the microstimulators, except for the tip of microelectrodes.

### Wireless coil fabrication

The Tx coil and the relay coil were fabricated with polyimide printed circuit boards and encapsulated with biomedical epoxy and LCP, respectively [51]. After attaching resonance capacitors (0.8 pF in parallel and 1.2 pF in series) to the Tx coil using solder paste (SMD4300SNL10, Chip Quik), we applied a minimal amount of biomedical epoxy to encapsulate the Tx coil. For the relay coil, we used thermoplastic LCP packaging to ensure the long-term safety of the coil. This packaging used multilayer LCP films combining high (CTZ-50, Kuraray) and low (CTF-25, CTF-50, Kuraray) melting temperature variants to improve interlayer adhesion and minimize defects. The thermocompression process using a heating press (model 4386, Carver Inc.) initially involved pressing films at 250 °C with a pressure of ∼ 250 kPa for 10 min to shape the top layer, then pressing at 284 °C with ∼ 450 kPa for 30 min to seal the top and bottom layers after aligning the coil in between. After fabrication, the hermetically packaged coil was subjected to an accelerated aging test at a controlled temperature of 120 °C for 3 days to confirm that the coil exhibited no changes in resonance properties. Any infiltration of water lowers the resonance frequency and the quality factor of the coil, which was identified through S-parameter measurements using the network analyzer (E5071B, Agilent). After the implantation of the relay coil in the rat, we checked its resonance using a portable network analyzer (MS2036A, Anritsu) as shown in Supplementary Figure 10.

### RF hardware for downlink command generation and uplink monitoring

We performed experiments with the stimulator chips using an SDR (the Model ‘Raptor’ by Rincon Research). For benchtop characterization, we used a single-turn printed circuit board including the Tx coil (9 mm × 9 mm) and wire-bonded chip pads to the board, connecting chips with a load resistor (14 kOhm) or electrodes inside the saline (Supplementary Figure 3). The SDR, using both transceiver chips (AD9361, Analog Devices) and Zynq SoCs, generated an RF downlink at 898 MHz, which was further amplified by an RF power amplifier (ADL5605-EVALZ, Analog Devices). The downlink ASK-PWM waveform was programmed on MATLAB, transferred to SDR, and imported to the field programmable gate array of the SDR using the IIO Oscilloscope library, which is controlled by MATLAB with the SSH protocol. The downlink signal is delivered to the Tx coil through an RF surface acoustic wave duplexer (D5DA942M5K2S2, Taiyo Yuden), which also isolates the backscattered signals from the microstimulators. The SDR amplifies the backscattered signal, downconverts it from 928 MHz (backscattering band) to DC, and finally digitizes the signal at 30 MSa/s (12-bit). The digitized data was then transferred to a personal computer for address and clock data extraction through BPSK demodulation and for calculating backscattering signal amplitude (computation performed in MATLAB).

### Animal surgery and device implantation

The microstimulators were sterilized using ethylene oxide before the implantation. We performed surgeries on two Long Evans male rats (500–700 g) for acute evaluation and chronic experiments. After shaving the animal’s hair, we secured it to a stereotaxic frame and sterilized the head skin. We then made a sagittal incision up to 5 cm long in the skin over the skull, followed by a craniotomy using a precision surgical dental drill. For the acute evaluation, we performed a craniotomy on the left hemisphere and implanted six microstimulators and a 32-channel penetrating multielectrode array from Blackrock Neurotech. While delivering the stimulations through wireless microchips, the explorer Nomad system (Nomad, Ripple Neuromed, sampling rate of 30 KSa/s) captured the neural responses from the electrode array. Recorded signals were first filtered with a 60 Hz notch filter to remove environmental noise, and then filtered again with a bandpass filter with a low cutoff of 60 Hz and a high cutoff of 2000 Hz. This was done to capture both the post-stimulus evoked LFP responses and the stimulation/downlink artifacts. During the initial surgery, the animal was anesthetized with 2-3% isoflurane but once the implantation was over, we switched anesthesia to ketamine/xylazine mixture (80 mg/kg and 5 mg/kg, respectively) to monitor the neural response better.

In the case of chronic surgery, we performed a bilateral craniotomy and manually implanted fifteen microstimulators on each side of the hemisphere. For the implantation, we adjusted the location of the implant to avoid the blood vessels. After the implantation, the open craniotomy was covered with the LCP-packaged relay coil, and we applied surgical PDMS first at the edge of the craniotomy and then covered the relay coil edge with dental cement. Lastly, sutures closed the surgical area, and the animals were allowed to recover from anesthesia before being transferred to the cage facility for post-surgical monitoring. Animals were continuously monitored throughout the studies and euthanized with pentobarbital sodium according to approved methods upon study conclusion. The implantation surgery for the chronic experiment was conducted under 1-3% isoflurane. All research protocols were approved and monitored by the Brown University Institutional Animal Care and Use Committee. Additionally, all research activities were conducted in compliance with relevant NIH guidelines and regulations.

### Microscale CT imaging

Micro-CT imaging was performed on a postmortem skull preserved in 4% paraformaldehyde using a SkyScan 1276, Bruker. The scan setting included an effective image pixel size of 10 μm, 1 × 1 binning, with a total of 1800 projection views made in a full 360-degree scan, along with an exposure time of 1250 ms. A peak source voltage of 100 kV and a current of 200 μA were used. The X-ray beam was filtered with a 0.5 mm thick aluminum filter and a 0.03 mm copper filter. Reconstruction was done using a Hamming filter (alpha = 0.54) processed by the NRecon software (v. 2.0.0.5). We collected micro-CT imaging to verify the positions of the microstimulators and evaluate the status of the relay coil throughout this period.

For that, the specimen was decapitated after being fixed in 4% paraformaldehyde 105 days after the implantation. Postmortem pictures are shown in Supplementary Figure 14.

### Design of the behavioral task and the apparatus

We built a custom enclosure with a motorized swivel using a 5 V stepper motor (28BYJ-48) and the driver board (ULN2003) controlled by Arduino. Since thin SMA cable (36 AWG, Alpha wire) cannot generate sufficient force for any shaft rotation, we manually controlled with rotation of the SMA connection using buttons, which are connected to a rotatory joint SMA adaptor (CR4794, Centric RF). On one side of the enclosure, we installed two levers for the detection and report task. The lever is made with photosensitive resin using a 3D printer (Mars 3, Elegoo) and connected to a snap-action switch. Levers were connected to Arduino boards for detection. We used MATLAB to collect information from the Arduino board during the behavioral task as well as to generate a downlink for intracortical microstimulation. For the lever task, once stimulation was delivered, the animal waited until it heard the report cue (1 kHz, 1s, generated by MATLAB), and it pressed the lever to report within 5 seconds. When the animal pressed the correct lever, the liquid reward, 10% diluted condensed milk (California farms sweetened condensed milk), was given while the total amount of the liquid was limited to 20 ml per day. If the animal pressed the lever when it was not reporting period, the initiation of the trial was delayed for 3 seconds to prevent the animal from randomly pressing the buttons.

### Animal tracking and movement analysis

When stimulating the motor cortex or tracking the movement of the animal in the enclosure, we used two Hero 7 Black (GoPro 1080p HD 1920 × 1080, 30 or 60 FPS) cameras mounted to the top and side of the enclosure. Two cameras share the audio input (random noise generated from MATLAB) to synchronize the frame as illustrated in Supplementary Figure 8. This sound was also captured by the oscilloscope (Tektronix MSO 5204) which monitors RF downlink in parallel. Using the oscilloscope and camera recording, we could precisely find the stimulation and downlink timing in the video. Tracking of the animal head’s or whisker’s position for either top-view or side-view videos was performed using the DeepLabCut package (Version 2.3.8) [52]. Body features, such as ear, nose, whisker base and tip, were chosen for the tracking. The training was conducted separately for each camera view using a GPU (NVIDIA GeForce GTX 1070) with 200,000 iterations. The tracking session results were exported to Excel format, which included the X and Y coordinates along with the confidence value (Likelihood) for each data point. The data points with a confidence value greater than 99% were used for head tracking and analysis for head velocity, distance traveled, and head angular velocity. Meanwhile, coordinates with a confidence value above 70% were selected to capture the fast oscillatory movements of the whiskers with manual error correction applied.

## Statistical analysis

Results in this work are presented as mean values ± s.d. (standard deviation) or mean values ± s.e.m. (standard error of the mean), which are described in the corresponding figure legends. Collected data are analyzed by MATLAB R2022a with custom code.

## Acknowledgments

The authors acknowledge F. Laiwalla at Brown University for her previous work on ASIC circuit block design, D. Durfee for his insights on circuits and medical devices, and J. Rosenstein for sharing his expertise and laboratory in the post-processing of chips. We also thank Y.-K. Song at Seoul National University for guidance in ASIC and hardware design. We have greatly benefited from the insight of our colleagues across multiple fields from microelectronics to brain sciences and clinical neurology: P. Asbeck at UCSD and S. Cash at Harvard Medical School. This research was supported by private gifts and NIH Award 1S10OD025181 (Brown University for computational resources).

## Contributions

J.L. and A.N. conceived the project. A.N., J.L., and A.-H.L. designed the animal experiments and methodology. J.L. and A.-H.L. designed and characterized the wireless ASICs and performed all experiments. J.L., A.-H.L., and A.N. wrote the manuscript. All authors contributed to the data analysis and provided feedback on the manuscript.

## Data availability

The data that supports the findings of this study are available from the corresponding author upon reasonable request.

## Code availability

Custom-developed MATLAB code is available from the corresponding author upon reasonable request.

