## Supplementary for "A Wireless Network of Implantable Microstimulators for Addressable and Patterned Electrical Brain Stimulation"

**Contents**

**Supplementary Note 1. Advantages of wireless daisy-chain communication for fast multichannel neural actuation.**

**Supplementary Note 2. Deploying amplitude shift keying pulse width modulation (ASK-PWM) and demodulation techniques.**

**Supplementary Note 3. Daisy-chain detector and microstimulator controller**

**Supplementary Note 4. Other on-chip supporting circuits**

**Supplementary Table 1. Comparison between the proposed microstimulator and other state-of-the-art stimulators.**

**Supplementary Table 2. Seven types of three-bit commands.**

**Supplementary Table 3. P-values for data comparison.**

**Supplementary Video 1: Head movement induced by a single microstimulator.**

**Supplementary Video 2: Whisker oscillatory movement induced by multiple microstimulators.**

**Supplementary Video 3: Lever press task for stimulation detection.**

**Supplementary Figure 1. Intracortical microelectrode fabrication and integration with stimulator ASIC.**

**Supplementary Figure 2. Current driver circuit configuration for biphasic stimulation.**

**Supplementary Figure 3. Benchtop setup for measuring current injection from the microstimulator.**

**Supplementary Figure 4. Coexistence of stimulation artifacts and LFP responses.**

**Supplementary Figure 5. Location of implanted microstimulators post-surgery and micro-CT scans.**

**Supplementary Figure 6. Relay and Tx coils for rodent experiments.**

**Supplementary Figure 7. Instrumentation on enclosure for chronic rodent studies.**

**Supplementary Figure 8. Schematic diagram of RF hardware for generation of downlink commands and uplink (backscattered) signal collection.**

**Supplementary Figure 9. Cortical location of selected microstimulation sites in the lever task.**

**Supplementary Figure 10. S-parameter measurements to quantify the resonance properties of the implanted relay coil with the animal temporally anesthetized using isoflurane.**

**Supplementary Figure 11. Location of the animal in the various coordinate planes within the enclosure and corresponding threshold measurements of microchip activation at each location.**

**Supplementary Figure 12. Images of video capture illustrating various activities of the freely moving rat and the corresponding RF incident power thresholds for microchip activation.**

**Supplementary Figure 13. Simulation of specific absorption rate (SAR) and its dependency on the duty cycle of RF transmission.**

**Supplementary Figure 14. Photographs of the 30 implanted microchips and the relay coil (postmortem).**

**Supplementary Note 1. Advantages of wireless daisy-chain communication for fast multichannel neural actuation**

Our daisy-chain wireless RF protocol allows each microchip to be programmed for specific pulsed current parameters by a 3-bit command-and-control downlink signal within 3 µs. Thus, for example, if the full stimulator network consists of 1000 implantable microchips, the command-and-control across the population can be accomplished in 3 ms (plus an additional 4 µs for SYNC). In the case of macroscale electronic devices, alternate RF communication approaches can be used for remote actuation; however, the distributed microchip system presents unique challenges. Each device is an ultra-miniaturized integrated circuit with a unique on-chip address and must operate at extremely low power. Thus, for example, accurate clock generation is not feasible as such circuits consume power and real estate. The flip side is having a network with variable clock frequencies across the chip population. As we show in the main and supplementary text, implementing a daisy-chain protocol with ASK-PWM (see Supplementary Note 2 next) offers a power efficient, low latency, and scalable solution that is also tolerant of variable clock frequencies. An important benefit from the short-pulse nature of the downlink commands in the daisy-chain method is the low average RF power, which complies with the SAR safety limits. We note that the method is not limited to electrical microsimulation and can benefit any type of downlink scheme where multiple downstream devices need to be programmed in a short period of time.

We had earlier considered an alternative approach to achieve comparable effectiveness in downlink communication via a call-and-response time-division multiple access (TDMA) [1]. In this method, the downlink includes the address of a specific target chip —10 bits in our case — followed by a 3-bit command for stimulation, thereby consuming 13 µs to communicate with a single chip (1 Mbps). In addition, with call-and-response TDMA, if only one SYNC is followed by address and command sequences for all chips, any combination of subsequent address and command bits can inadvertently trigger chips other than the intended target to deliver the current stimulus. To prevent false stimulation in chips, each downlink call must include the SYNC, address, and command only for a single target chip, with the SYNC taking an additional 4 µs. As a result, a total of 17 µs time window is required to program one chip, 17 ms for a population of 1000 chips, which is more than 6 times slower than the daisy-chain method. Additionally, the call-and-response TDMA is less energy efficient than the daisy-chain. This is because, in call-and-response TDMA, each chip must frequently detect the SYNC and compare the incoming 10-bit address with its unique address, whereas, in the daisy-chain method, the detector only needs to wait and find the target 3-bit command.

**Supplementary Note 2. Deploying amplitude shift keying pulse width modulation (ASK-PWM) and demodulation techniques**

Since each microstimulator uses a free-floating oscillator for clock generation, we designed and deployed an ASK-PWM downlink modulation. Here, a Bit 0 consists of a short high-amplitude signal (333 ns) followed by a long low-amplitude signal (666 ns), while Bit 1 comprises a long high-amplitude signal (666 ns) followed by a short low-amplitude signal (333 ns) [1]. The on-chip oscillator has a nominal clock frequency of 30 MHz (clock period of 33 ns) and drives 'counter-high' and 'counter-low'. These circuits count the periods of high and low amplitudes of the incoming RF signal, respectively. When comparing the relative lengths of the periods of high and low amplitude, if the output from the 'counter-high' ($N_H$) exceeds that of 'counter-low' ($N_L$) the demodulated bit is 1; otherwise, the bit is 0. Therefore, in the ASK-PWM demodulation process, the relative ratio between the downlink rate and the clock frequency determines $N_H$ and $N_L$. However, since the clock frequency of the specific chip is fixed, we varied the downlink rate from 0.73 MHz to 1.26 MHz instead with the results shown in Figs. 2e and 2f. In this case, $N_H$ and $N_L$ will be inversely proportional to the downlink data rate but this should not affect the demodulated downlink bits. As expected, the ASK-PWM demodulator demonstrated wide tolerance to variations in the downlink data rate and correctly responded by generating stimulation at the exact timing. This result explains how the population of stimulators reliably responded to the downlink even with the variance in clock frequency ranging from 28 to 32 MHz in subsequent experiments. To drive the ASK-PWM downlink, we designed the Tx waveform by setting the amplitude of the 'high' status to be 25% higher than the nominal status and the 'low' status to be 25% lower than the nominal status. The final demodulated downlink bit had a data rate of 1 Mbps.

**Supplementary Note 3. Daisy-chain detector and microstimulator controller**

The daisy-chain detector on each microchip receives downlink bits, a chip-specific address, and a 10 MHz clock. This detector also receives a separate input signal from the ASK-PWM circuit, which detects the unique SYNC waveform in the downlink RF signal. This SYNC waveform consists of a very long high-amplitude signal (1666 ns), a short low-amplitude signal (333 ns), another very long low-amplitude signal (1666 ns), and a short high-amplitude signal (333 ns). Therefore, no signals for the downlink command bits can be misinterpreted as the SYNC waveform, considerably improving the robustness of the downlink. Once the SYNC waveform is detected, the daisy-chain detector monitors the following demodulated downlink bits: the first three bits are dedicated to Chip 0, the next three bits to Chip 1, and so forth. If the chip address is 30, the daisy-chain decoder will select the bits from the 91st to the 93rd after the SYNC, for example. Therefore, the chip address determines the specific timeslot for each chip, which is a laser-written address programmed after CMOS fabrication (by ablation) as described in references [1, 2]. In the present stimulator ASIC, there are 10-bit metal fuses for addressing thereby allowing up to 1024 usable addresses. After finding a 3-bit command for the target chip, the daisy-chain detector feeds it to the stimulation controller, which then dictates the amplitude and pulse width of the current stimulus based on preprogrammed parameters.

**Supplementary Note 4. Other on-chip supporting circuits**

The stimulator ASIC features a 3-stage cross-coupled rectifier for efficient RF-DC conversion in near-field energy harvesting. The rectifier also serves as an envelope detector; thus, the output voltage of the rectifier (Vrect) reflects the amplitude changes in the downlink as shown in Fig. 1d. The envelope signal is processed in the ASK-PWM demodulator, which compares Vrect with $V_{DD}$. $V_{DD}$ is the low-pass filtered signal of Vrect. By doing so, the demodulator generates binary digital data where a high amplitude results in an output bit of 1, and a low amplitude results in an output bit of 0. This period of bit 1 and bit 0 is compared by the counter-high and counter-low for the demodulation as described in Supplementary Note 2 above. Unregulated $V_{DD}$ provides the power to power-on reset (POR), ASK-PWM demodulator, BPSK modulator, daisy-chain detector, controller, and stimulation driver. Since the unregulated $V_{DD}$ can increase beyond the nominal voltage of 1 V, we used an over-voltage protection (OVP) diode to limit the voltage increase. The POR circuit triggers the BPSK modulator to transmit the uplink at the nominal data rate of 10 Mbps, which includes a total of 960 bits (repeating 32-bit predefined LFSR and 10-bit address bits). Backscattering is initiated by toggling the transistor connected to the capacitor which affects the coil's resonance. The BPSK modulator uses the clock frequency of the freely running on-chip oscillator, nominally at 30 MHz. Thus, the backscattered signals carry information about the address and the clock frequency. Since the clock frequency depends on $V_{DD}$, it serves as an indicator of the on-chip power level (Fig. 2a). For a population of microchips, backscattering can be used to determine whether all chips are switched on as shown in Fig. 5c. Whenever a chip is switched on, it begins to generate its backscattering uplink. However, during the uplink transmission, the chip responds less reliably to the downlink because the toggling of the transistor also affects the on-chip power. As a remedy, we can interrupt the backscattering by turning the chip on and off in a very short period of time, allowing it to reliably respond to the downlink command - without waiting for the uplink to finish. As for the current driver, we used a programmable current source architecture with associated switches to generate biphasic electrical stimulation between two differential pads, as shown below in Supplementary Figure 2. We set the upper limit for the current stimulation at 60 µA due to its application in the intracortical area; however, it can be scaled beyond this limit by modifying the current source.

| **Table S1\| Comparison between the proposed microstimulator and other state-of-the-art stimulators** | | | | | | |
|---|---|---|---|---|---|---|
| | [3] | [4] | [5] | [6] | [7] | This work |
| Number of node(s) | 1 | 1 | 1 | 1 | 1 | 30 (Up to 1000) |
| Link | Ultrasonic | Inductive | Magnetoelectric | Mid-field | Inductive | Inductive |
| $f_{carrier}$ (MHz) | 1.85 | 1300 | 1.25 | 1600 | 13.56 | 898 |
| Uplink | Backscattering (OOK) | None | None | None | 433 MHz OOK | Backscattering (BPSK) |
| Downlink | - | - | 4.6 kbps | - | - | 1 Mbps |
| Process (nm) | 65 | 130 | 180 | - | 130 | 65 |
| Supply voltage (V) | Up to 2.5 | 1.2 | 1.8 | 2 | 2 | 0.8 |
| Stimulation start | Stimulation command | Received power | Stimulation command | - | Stimulation command | Stimulation command |
| Max. current level (µA) | 400 | 38 | - | - | 1860 | 120 peak-to-peak |
| Charge balance | Passive recharge | Biphasic | Biphasic | - | 3-bit biphasic correction | Biphasic |
| Animal model | Acute/Rat sciatic nerve | Acute/Rat sciatic nerve | Acute/Pig femoral nerve | Acute/ Rabbit heart | Rat sciatic nerve | Chronic / Rat Cortex |
| Fully implanted | Yes | No | Yes | Yes | Yes | Yes |
| In-vivo model distance $_{Tx-Rx}$ (mm) | 18 | - | 15 | 45 | - | 3-5 |
| IC Area ($mm^2$) | 0.56 | 0.09 | 0.8 | - | 22 | 0.25-0.09 |
| Energy harvester | Piezoelectric | On-chip Coil | Magnetoelectric film | Discrete coil | Discrete coil | On-chip coil |
| Electrodes | PEDOT planar | Stainless steel disk on cuff | Microwire | Microwire | Cuff | Tungsten microwire |
| Implant encapsulation | Parylene | - | Epoxy | Epoxy | Epoxy and PDMS | Epoxy and parylene |

| Table S2\| Seven types of three-bits commands | | |
|---|---|---|
| **No. Command** | **3-bits** | **Nominal current amplitude and pulse width** |
| Command 1 | 001 | 20 μA, 100 μs per phase |
| Command 2 | 010 | 40 μA, 100 μs per phase |
| Command 3 | 011 | 60 μA, 100 μs per phase |
| Command 4 | 100 | 20 μA, 500 μs per phase |
| Command 5 | 101 | 40 μA, 500 μs per phase |
| Command 6 | 110 | 60 μA, 500 μs per phase |
| Command 7 | 111 | 100 Hz Burst, 60 μA, 500 μs per phase, continuous until new SYNC detected |

| Table S3\| P-values for data comparison (Figure 3h, Electrode 1) | | | | |
|---|---|---|---|---|
| | **C1, Command 3** | **C1/2/3, Command 3** | **C1, Command 6** | **C1/2/3, Command 6** |
| **C1, Command 3** | 1 | 0.00597 | 0.011 | $1.65 \times 10^{-7}$ |
| **C1/2/3, Command 3** | 0.00597 | 1 | 0.412 | $3.43 \times 10^{-5}$ |
| **C1, Command 6** | 0.011 | 0.412 | 1 | $7 \times 10^{-5}$ |
| **C1/2/3, Command 6** | $1.65 \times 10^{-7}$ | $3.43 \times 10^{-5}$ | $7 \times 10^{-5}$ | 1 |

A p-value $< 0.05$ indicates that the differences in the amplitudes of LFP responses between two different conditions are statistically significant (black background: statistically not significant, white background: statistically significant).

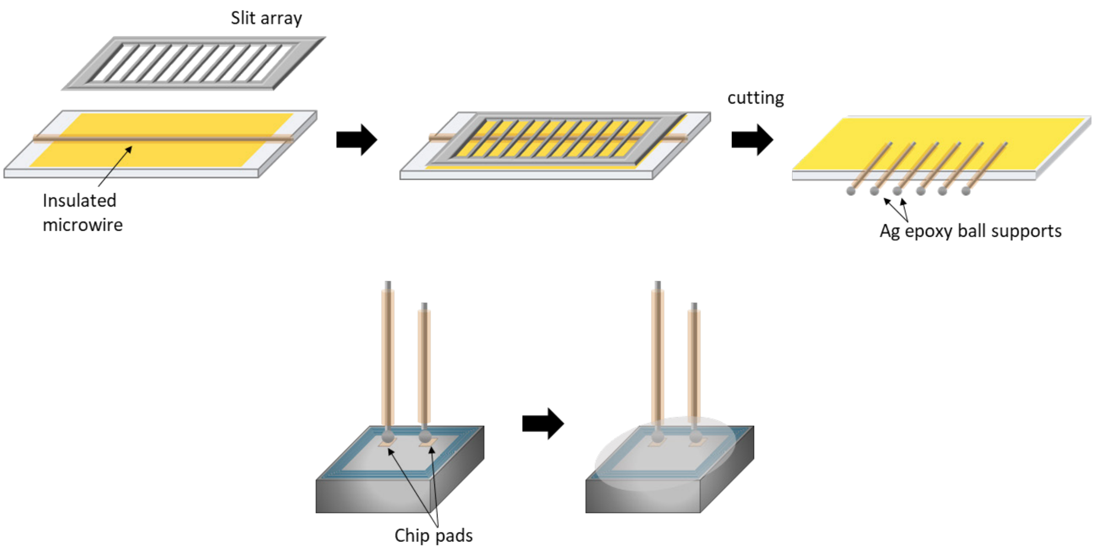


**Supplementary Figure 1. Intracortical microelectrode fabrication and integration with stimulator ASIC.** A long polyimide-insulated microwire is attached to a glass slide using double-sided Kapton tape, with a 3D printed resin slit array guide positioned on top for precision cutting. The wire is sectioned into desired lengths of 1 mm and 1.5 mm using a sharp blade. The edges of the cut wires are then stripped using a surgical blade and tiny metal ball supports are formed by applying a small amount of silver epoxy at the end of each microwire. Next, a pair of wires are aligned with the 60 µm size chip aluminum pads and secured using an additional small amount of silver epoxy, which is cured while maintaining perpendicular alignment to the chip surface. Once fully cured, a drop of biomedical-grade epoxy is applied around the silver epoxy to ensure mechanical stability and insulation. The microchips and electrodes, except for the tips of the electrodes, were then encapsulated by a thin film of parylene-C.

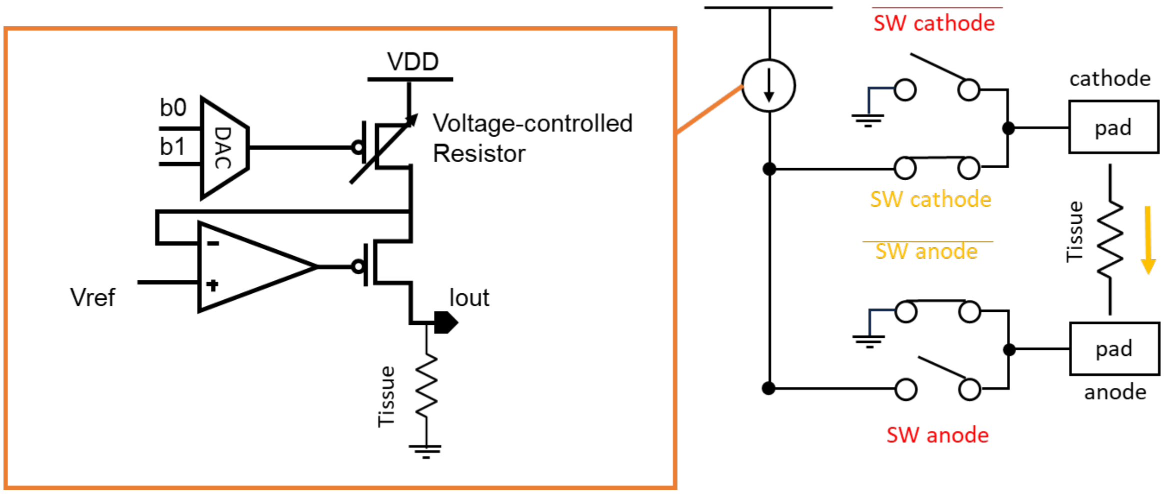


**Supplementary Figure 2. Current driver circuit configuration for biphasic stimulation.** Programming the parameters of current stimulus by a digital-to-analog converter (DAC) and a voltage-controlled resistor. Digital signals (b0, b1) are converted into specific voltage levels to modulate the current intensity (Iout) through the tissue. A feedback loop with a reference voltage (Vref) ensures consistent current delivery to tissue. The single current source efficiently injects current in alternating directions between the cathode and anode to provide the desired active charge-balanced biphasic stimulation.

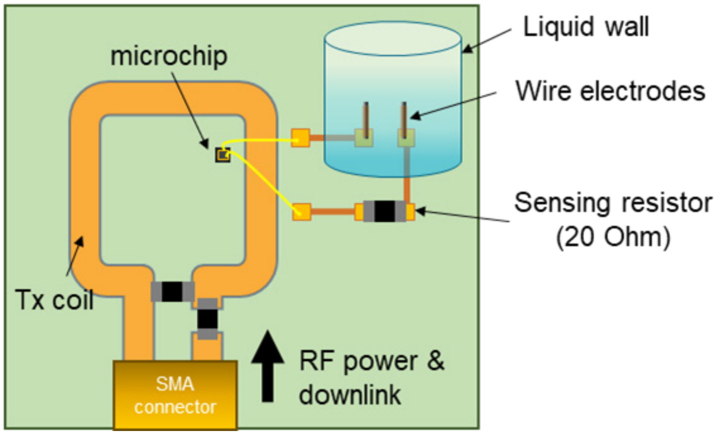


**Supplementary Figure 3. Benchtop setup for measuring current injection from the microstimulator.** Schematic of arrangement for measuring the current delivered through a pair of microwire electrodes by a wirelessly energized microstimulator. The schematic shows a transmitter (Tx) coil for wireless power transfer and associated components (20 Ω sensing resistor) wired on an FR4 printed circuit board (PCB) adjacent to a saline holding wall where the electrodes are located. The Tx coil is connected to an external RF hub through an SMA connector and cable.

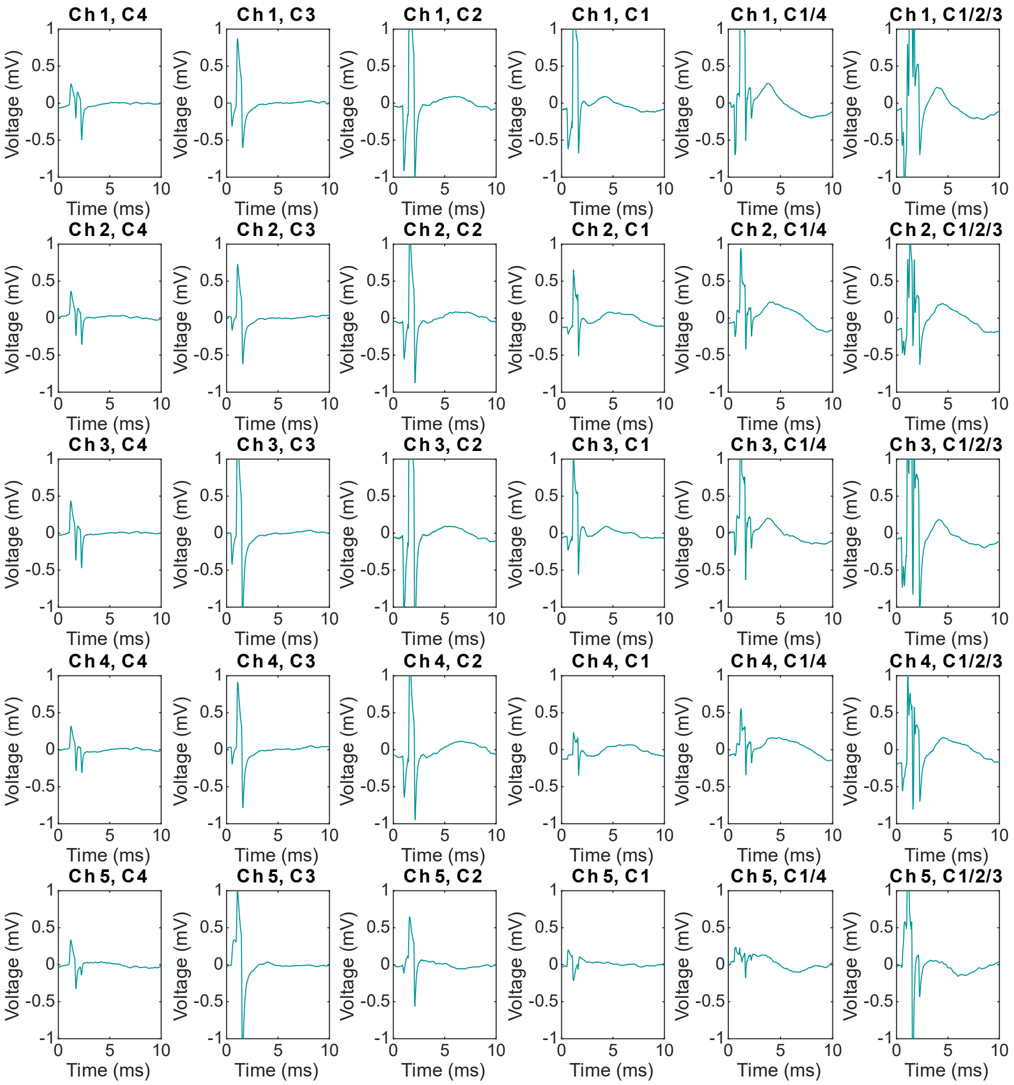


**Supplementary Figure 4. Coexistence of stimulation artifacts and LFP responses.** Ch 1-5 denote the recording electrodes of the multielectrode array (MEA) implanted adjacent to microchips to record neural responses. C1, C2, C3, and C4 are labels for the selected microstimulators. For example, 'C1/2/3' indicates that C1, C2, and C3 delivered the current stimulation simultaneously. Recorded signals are averaged over six trials.

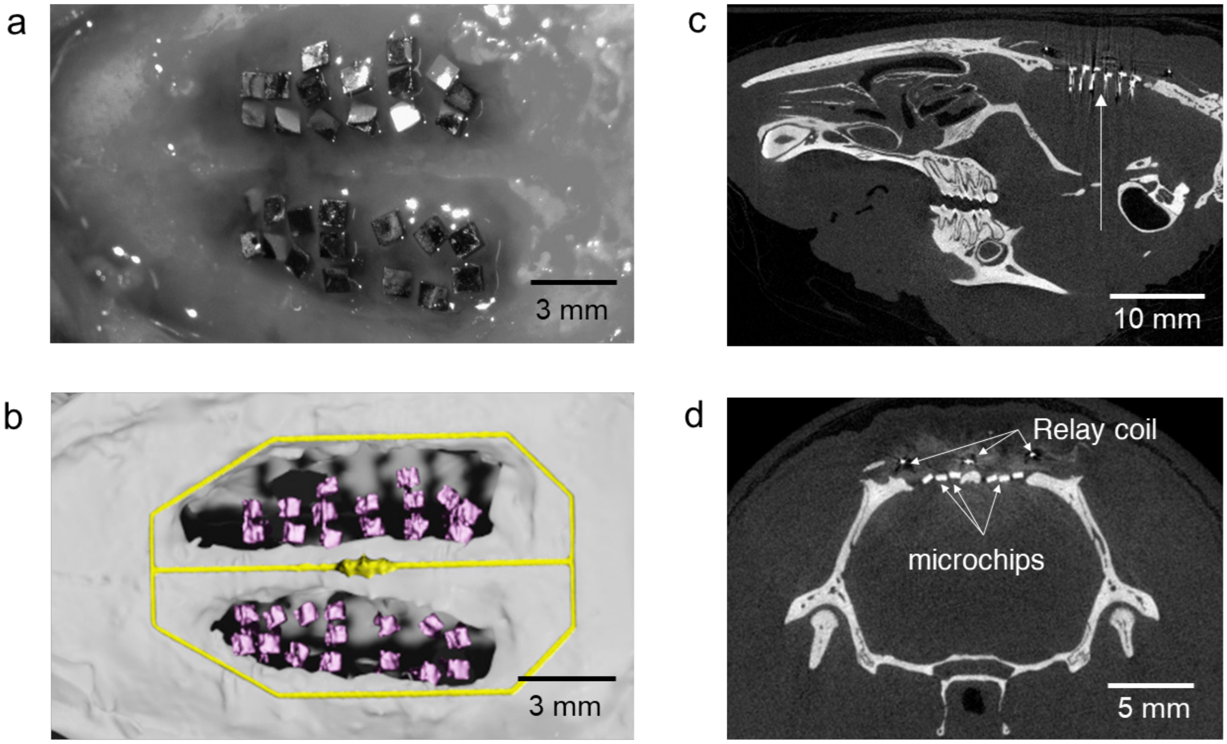


**Supplementary Figure 5. Location of implanted microstimulators post-surgery and micro-CT scans. a.** A photo showing 30 microchips implanted across the motor and sensory cortices of a rat, **b.** Image of a 3D reconstruction model generated from a sequence of postmortem micro-CT scans, **c.** Micro-CT scan in the sagittal plane showing microelectrodes penetrating the cortex. **d.** Micro-CT scan in the coronal plane showing the location of microstimulator chips and the relay coil. Postmortem pictures are shown in Supplementary Figure 14.

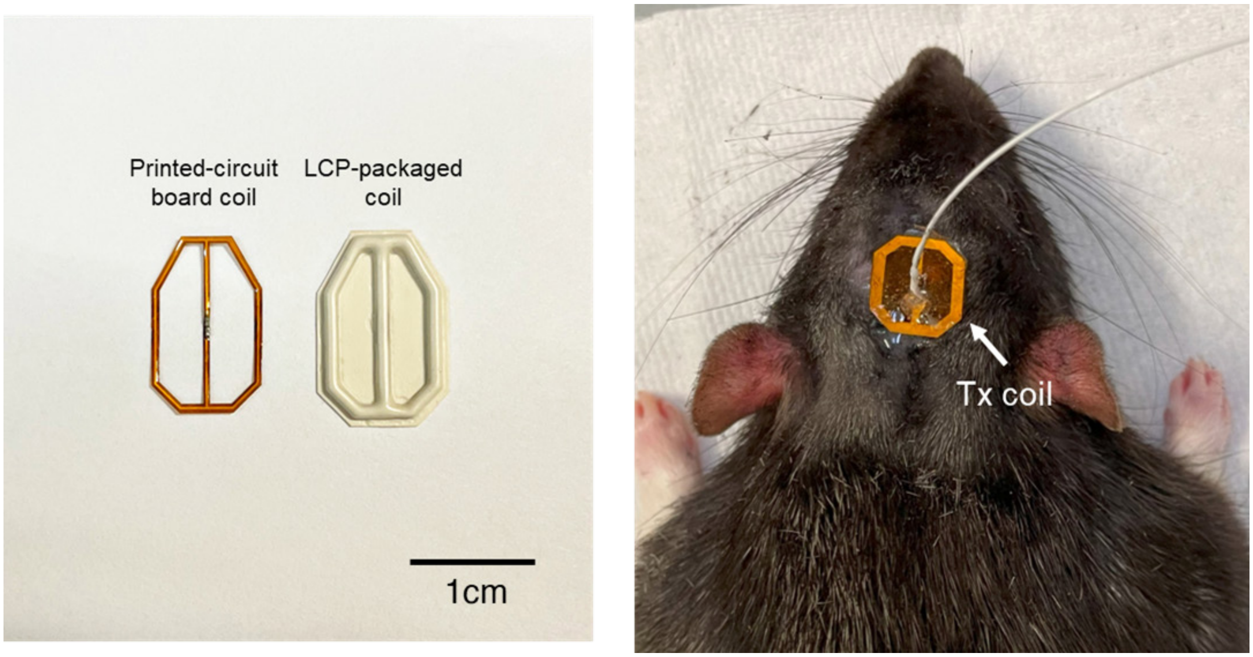


**Supplementary Figure 6. Relay and Tx coils for rodent experiments. a.** Printed circuit board coils, designed according to the size and shape of the rat's head and brain, with an integrated capacitor for resonance matching, shown before and after liquid crystal polymer (LCP) packaging. **b.** Tx coil mounted on the animal's head using surgical-grade Polydimethylsiloxane (PDMS).

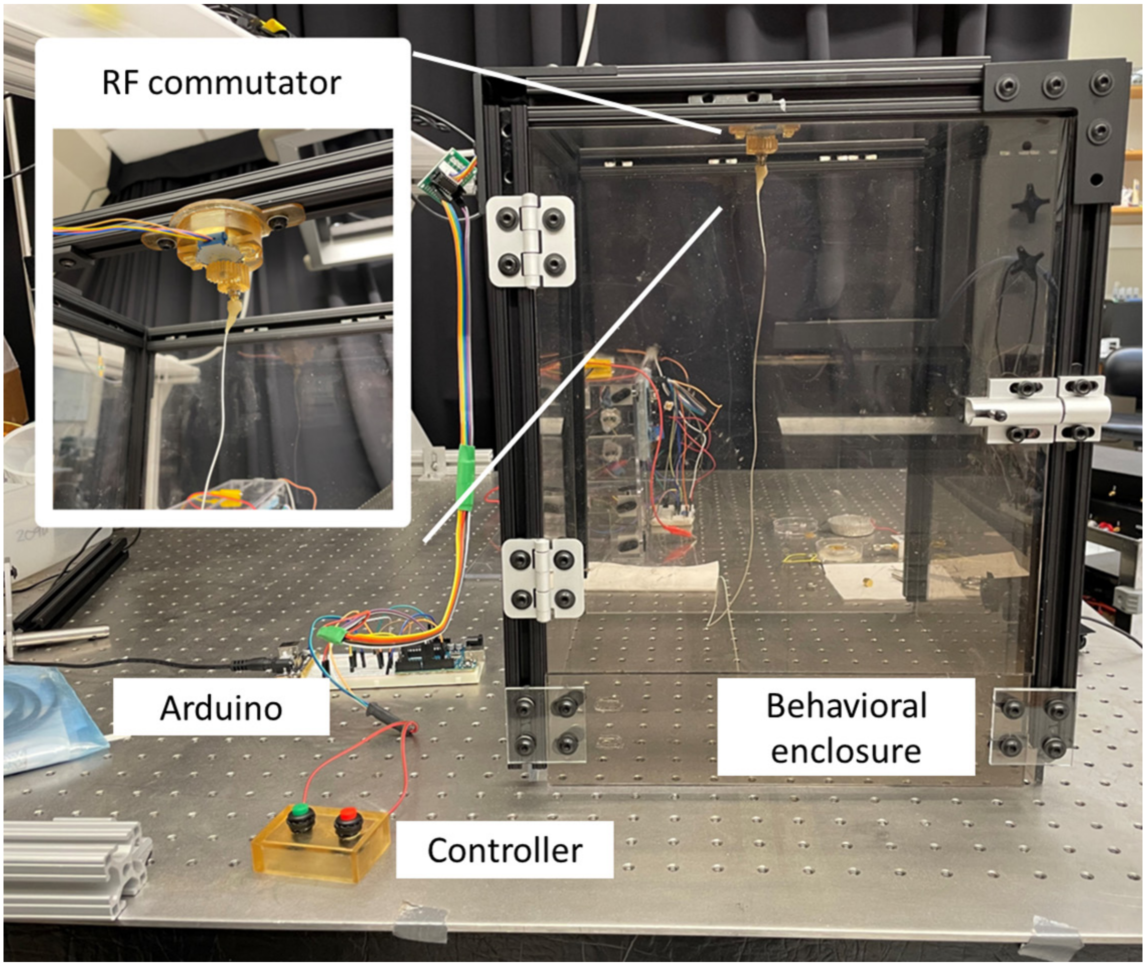


**Supplementary Figure 7. Instrumentation on enclosure for chronic rodent studies.** Enclosure for the freely moving animal (33 cm × 33 cm × 42 cm), with an RF commutator connector on top of the acrylic box. The RF commutator features a rotary SMC connector and a stepper motor, allowing the experimenter to untangle the micro-SMA cable as the animal moves and rotates. Since the micro-SMA cable is very thin (~1 mm), simple manual control of the stepper motor controller ('CONTROLLER') was sufficient.

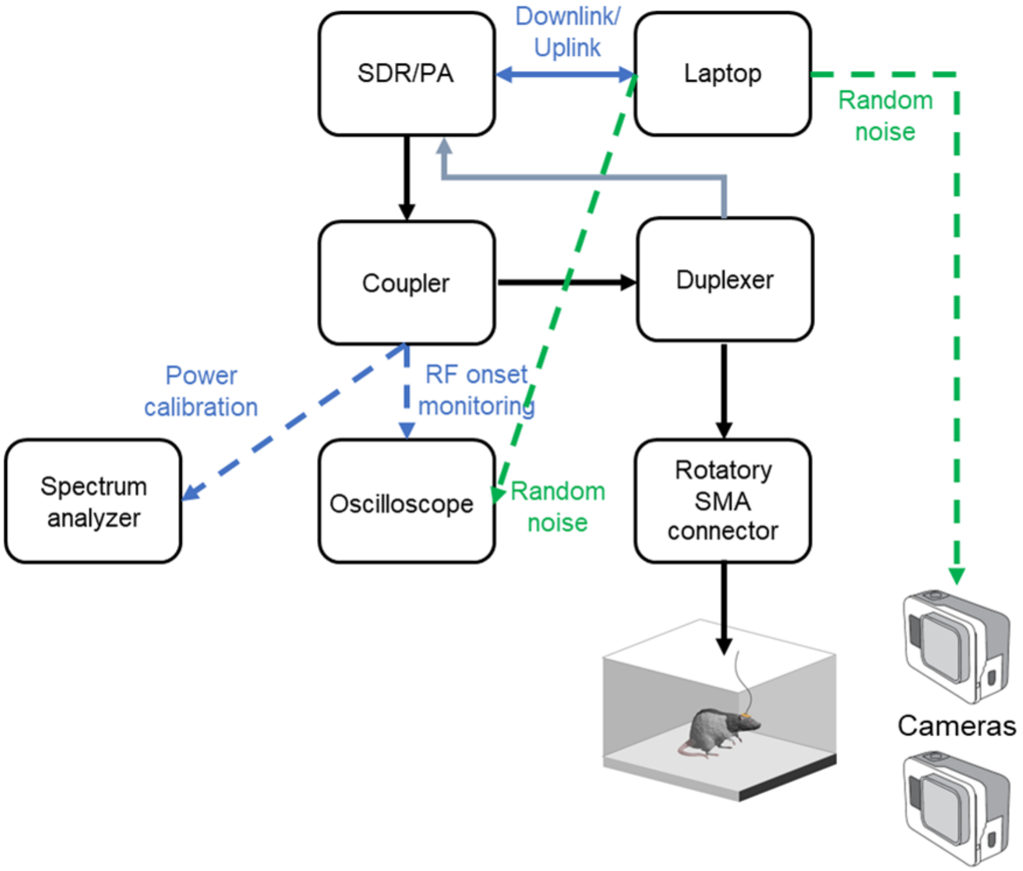


**Supplementary Figure 8. Schematic diagram of RF hardware for generation of downlink commands and uplink (backscattered) signal collection.** A software-defined radio (SDR) and power amplifier (PA) transmit the downlink signal while a directional coupler is used to monitor the timing and amplitude of the signal. The coupler is connected to a spectrum analyzer and an oscilloscope for monitoring purposes. A laptop controls the SDR using MATLAB and SSH communication, generating random noise as well. This noise is captured by the oscilloscope and cameras, enabling us to precisely identify when the downlink transmission occurs in the video.

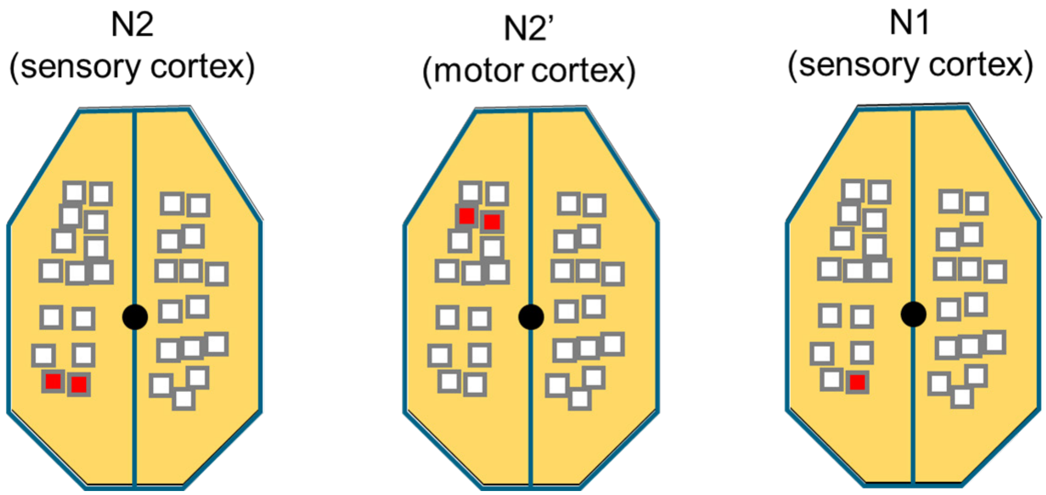


**Supplementary Figure 9. Cortical location of selected microstimulation sites in the lever task.** Filled red squares indicate those microchips which were wirelessly activated for the three sets of cortical stimuli (N2, N2', and N1).

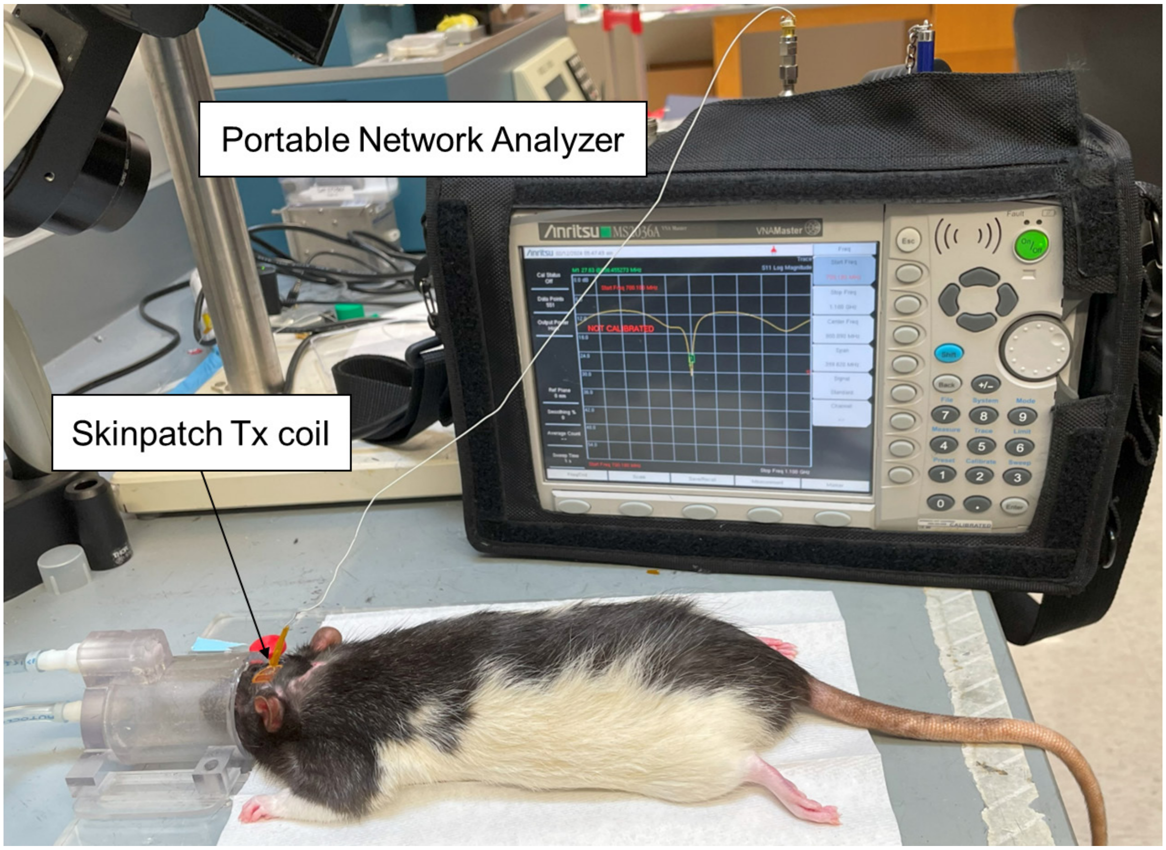


**Supplementary Figure 10. S-parameter measurements to quantify the resonance properties of the implanted relay coil with the animal temporally anesthetized using isoflurane.**

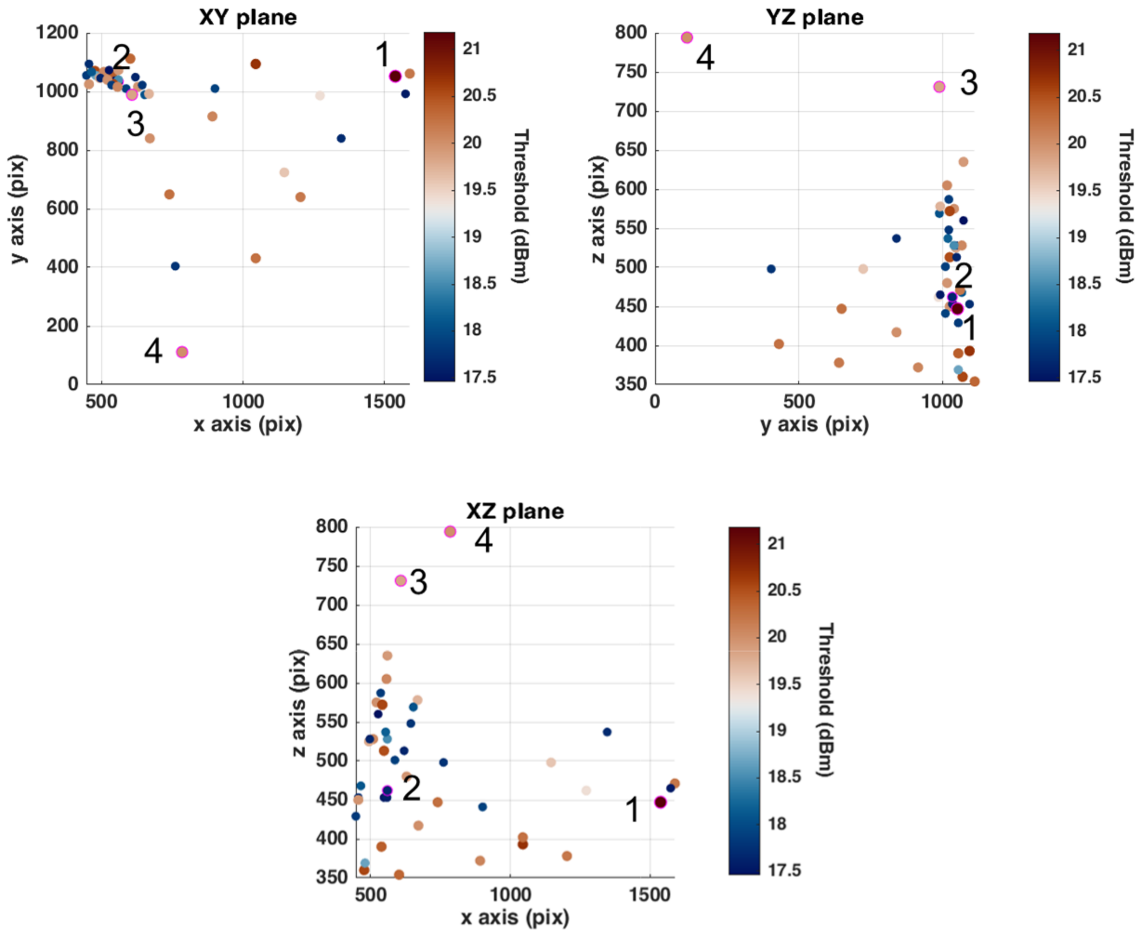


**Supplementary Figure 11. Location of the animal in the various coordinate planes within the enclosure and corresponding threshold measurements of microchip activation at each location.** Video frame capture for positions 1, 2, 3, and 4, respectively, are shown in the following Supplementary Figure 12.

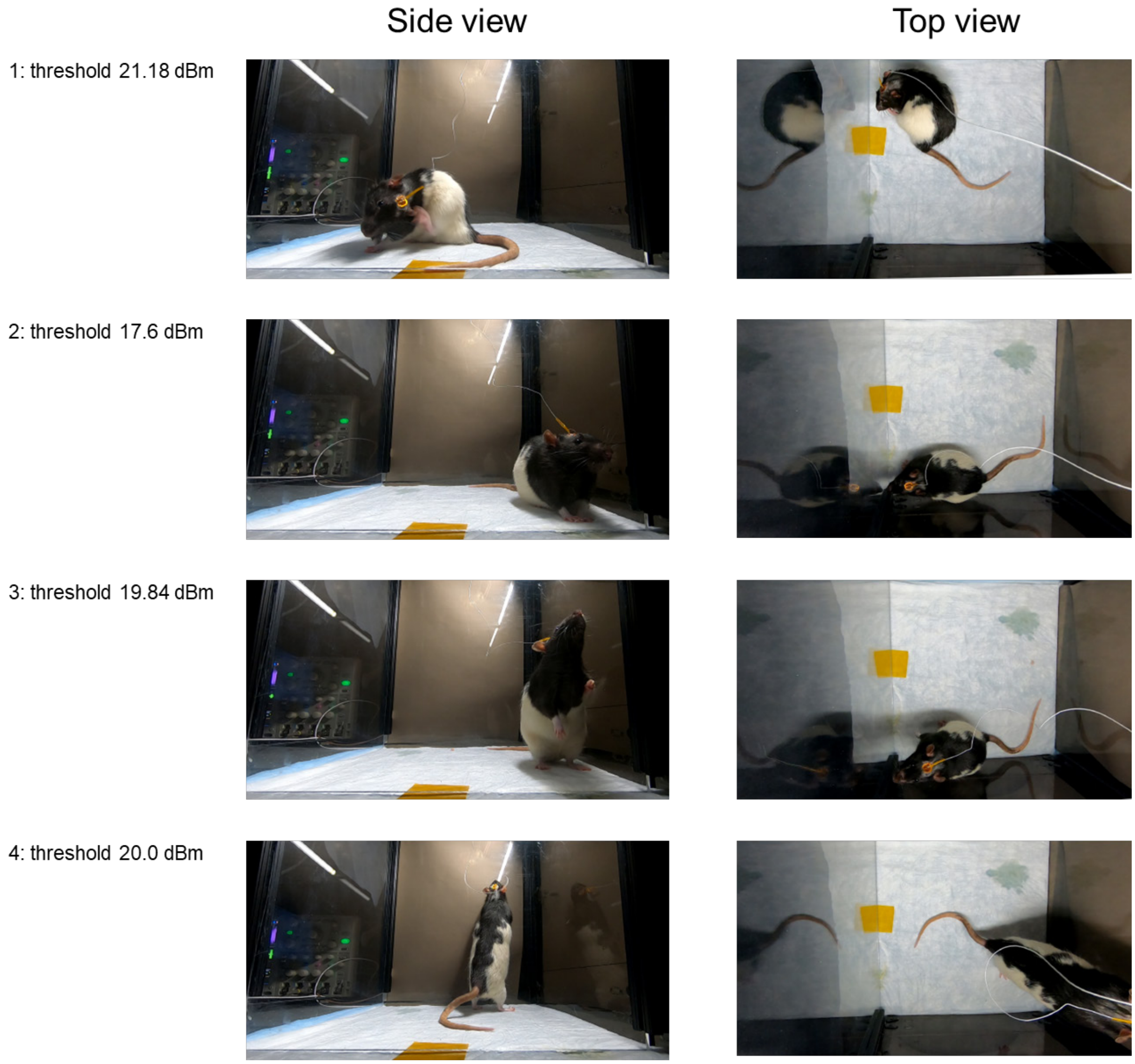


**Supplementary Figure 12. Images of video capture illustrating various activities of the freely moving rat and the corresponding RF incident power thresholds for microchip activation.** When the animal stays still, the threshold (dBm) is at its lowest. When the animal touches its coil and head directly with its paw, the threshold increases.

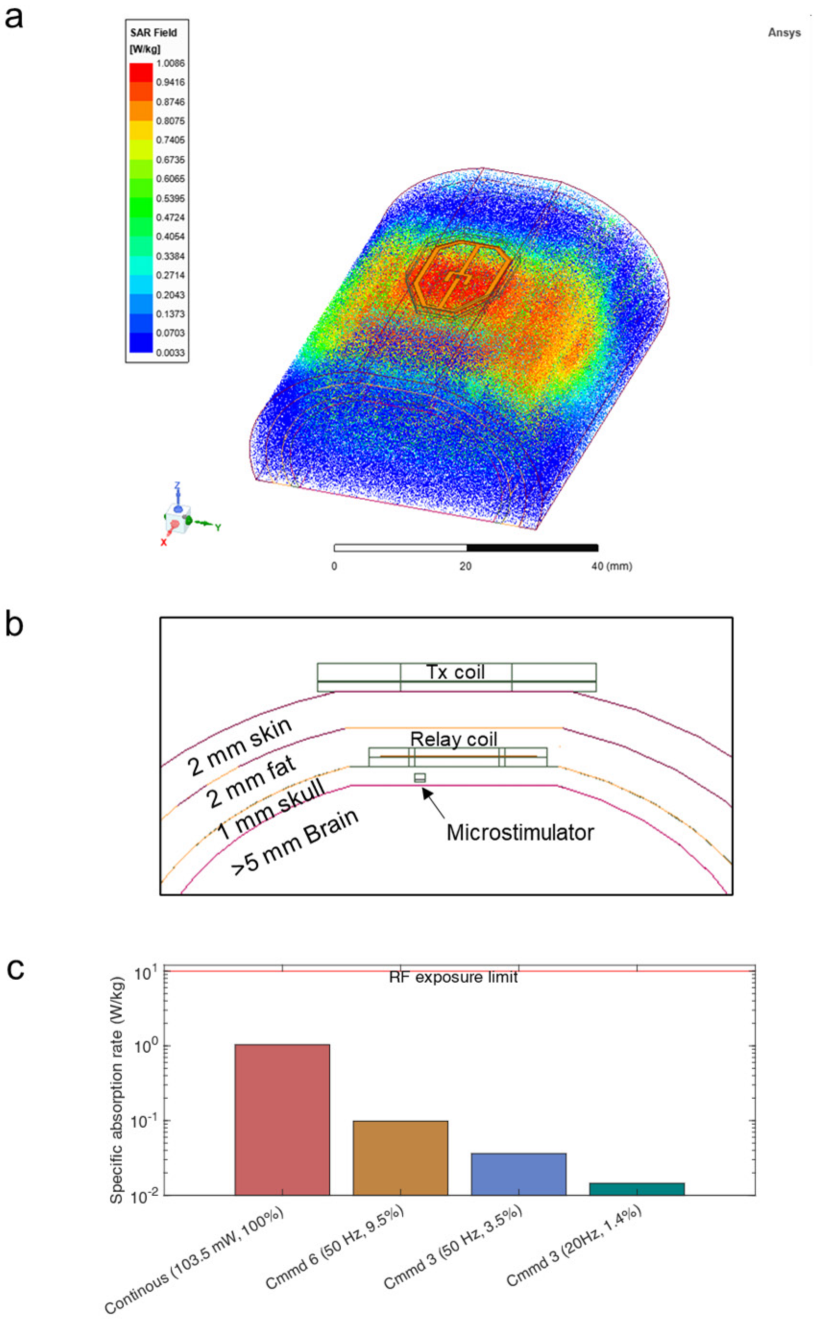


**Supplementary Figure 13. Simulation of specific absorption rate (SAR) and its dependency on the duty cycle of RF transmission. a.** Simulated SAR field with a 100 mW Tx emission and the 3-coil system specifically designed for the rodent model, as simulated on Ansys HFSS. Averaging SAR over 10-g tissue was done by following the procedure in IEC/IEEE 62704-4 [8]. **b.** The thickness of tissue layers used for the simulation (2 mm of skin, 2 mm of fat, 1 mm of skull, and more than 5 mm of brain tissue), **c.** SAR as a function of the duty cycle of RF transmission. Note that even for the lowest RF duty cycle a clear neural response to the current stimulus was measured. All SAR values in the figure lie well below regulatory limits (for humans).

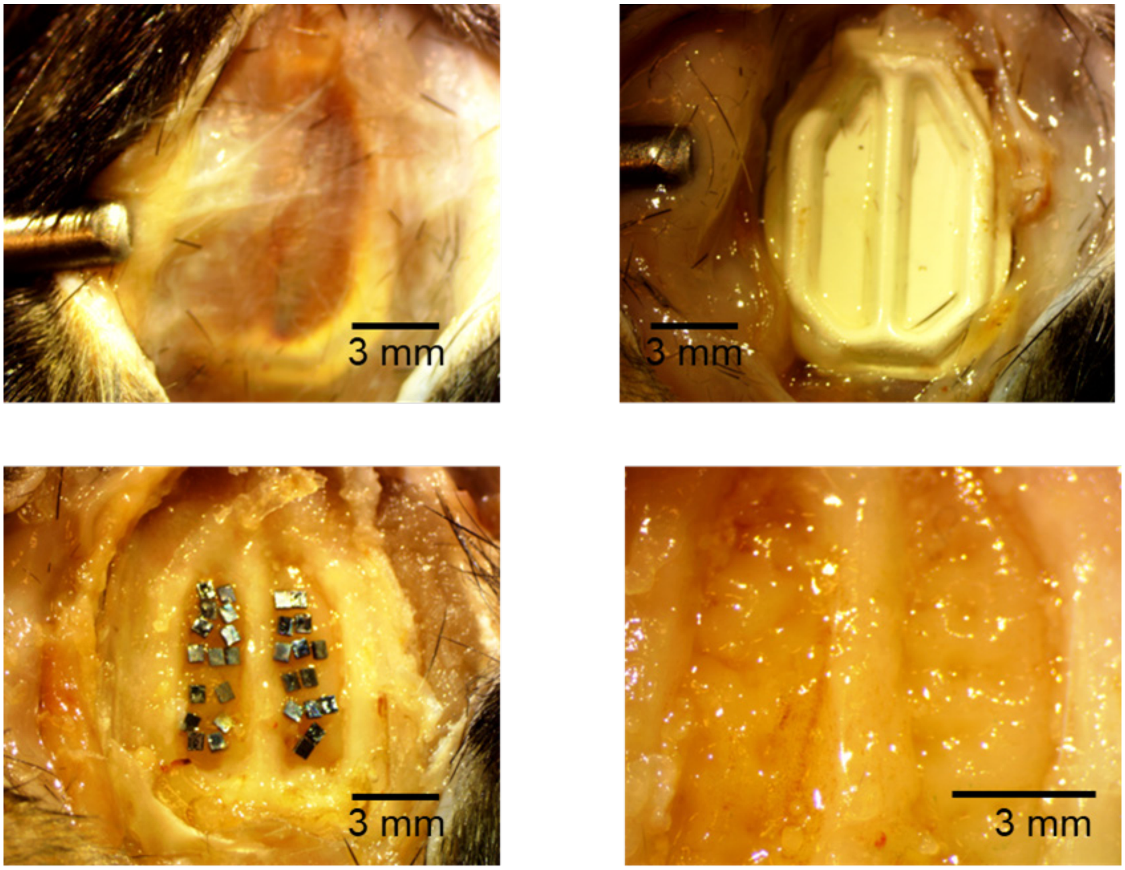


**Supplementary Figure 14. Photographs of the 30 implanted microchips and the relay coil (postmortem).** These images show that the relay coil is well integrated with the surrounding tissue and that the location of the microstimulators did not change observably over the period of the study.
